# Widespread ripples synchronize human cortical activity during sleep, waking, and memory recall

**DOI:** 10.1101/2021.02.23.432582

**Authors:** Charles W. Dickey, Ilya A. Verzhbinsky, Xi Jiang, Burke Q. Rosen, Sophie Kajfez, Brittany Stedelin, Jerry J. Shih, Sharona Ben-Haim, Ahmed M. Raslan, Emad N. Eskandar, Jorge Gonzalez-Martinez, Sydney S. Cash, Eric Halgren

## Abstract

Declarative memory encoding, consolidation, and retrieval require the integration of elements encoded in widespread cortical locations. The mechanism whereby such ‘binding’ of different components of mental events into unified representations occurs is unknown. The ‘binding-by-synchrony’ theory proposes that distributed encoding areas are bound by synchronous oscillations enabling enhanced communication. However, evidence for such oscillations is sparse. Brief high-frequency oscillations (‘ripples’) occur in the hippocampus and cortex, and help organize memory recall and consolidation. Here, using intracranial recordings in humans, we report that these ~70ms duration 90Hz ripples often couple (within ±500ms), co-occur (≥25ms overlap), and crucially, phase-lock (have consistent phase-lags) between widely distributed focal cortical locations during both sleep and waking, even between hemispheres. Cortical ripple co-occurrence is facilitated through activation across multiple sites, and phaselocking increases with more cortical sites co-rippling. Ripples in all cortical areas co-occur with hippocampal ripples but do not phase-lock with them, further suggesting that cortico-cortical synchrony is mediated by cortico-cortical connections. Ripple phase-lags vary across sleep nights, consistent with participation in different networks. During waking, we show that hippocampo-cortical and cortico-cortical co-ripples increase preceding successful delayed memory recall, when binding between the cue and response is essential. Ripples increase and phase-modulate unit firing, and co-ripples increase high-frequency correlations between areas, suggesting synchronized unit-spiking facilitating information exchange. Co-occurrence, phase-synchrony, and high-frequency correlation are maintained with little decrement over very long distances (25cm). Hippocampo-cortico-cortical co-ripples appear to possess the essential properties necessary to support binding-by-synchrony during memory retrieval, and perhaps generally in cognition.

**Significance Statement:** Different elements of a memory, or any mental event, are encoded in locations distributed across the cortex. A prominent hypothesis proposes that widespread networks are integrated with bursts of synchronized high-frequency oscillations called ‘ripples,’ but evidence is limited. Here, using recordings inside the human brain, we show that ripples occur simultaneously in multiple lobes in both cortical hemispheres, and the hippocampus, generally during sleep and waking, and especially during memory recall. Ripples phase-lock local cell firing, and phase-synchronize with little decay between locations separated by up to 25cm, enabling long-distance integration. Indeed, co-rippling sites have increased correlation of very high-frequency activity which reflects cell firing. Thus, ripples may help bind information across the cortex in memory and other mental events.

## Introduction

Ripples are brief high-frequency oscillations that have been well-studied in the rodent hippocampus during non-rapid eye movement sleep (NREM), when they mark the replay of events from the previous waking period, and are critical for memory consolidation in the cortex (1–4). Recently, ripples were found in rat association cortex but not primary sensory or motor cortices during sleep, with increased coupling to hippocampal ripples in sleep following learning (5). An earlier study reported ripples in waking and sleeping cat cortex, especially NREM (6). In humans, cortical ripples have recently been identified during waking, and were more frequently found in lateral temporal than rolandic cortex. Hippocampal sharpwave-ripple occurrence and ripple coupling between parahippocampal gyrus and temporal association cortex increase preceding memory recall in humans (7, 8), possibly facilitating replay of cortical neuron firing sequences established during encoding (9). In rats, ripples co-occur between hippocampus and ~1mm^2^ of parietal cortex in sleep following learning (5), in mice, ripples propagate from the hippocampus to retrosplenial cortex (10), and in cats, ripple co-occurrence is reportedly limited to short distances (6).

We recently reported, using human intracranial recordings, that ~70ms long ~90Hz ripples are ubiquitous in all regions of the cortex during NREM as well as waking (11). During waking, cortical ripples occur on local high frequency activity peaks. During sleep, cortical ripples typically occur on the cortical down-to-upstate transition, often with 10-16 Hz cortical sleep spindles, and local unit firing patterns consistent with generation by pyramidal-interneuron feedback. We found that cortical ripples group co-firing within the window of spike-timing-dependent plasticity. These findings are consistent with cortical ripples contributing to memory consolidation during NREM in humans.

While there is thus an emerging appreciation that hippocampal and cortical ripples have an important role in human and rodent memory, nothing is known of the network properties of cortical ripples. Specifically, it is not known if ripples co-occur or phase-synchronize between cortical sites, and if so, whether this is affected by distance or correlated with the reconstruction of declarative memories. These would be critical properties for cortical ripples to participate in the binding of different elements of memories that are represented in disparate cortical areas, the essence of hippocampus-dependent memory (12).

The binding of disparate elements of a memory is a specific case of a more general problem of how the various contents of a mental event are united into a single experience. Most often addressed is how different visual qualities of an object (e.g., color, shape, location, texture) are associated with each other (13), but the ‘binding problem’ generalizes to how the contents of awareness are unified in a single stream of consciousness (14). Modern accounts often rely on hierarchical and multimodal convergence. However, cortical processing is distributed, and it would be difficult to represent the combinatorial possibilities contained in all potential experiences with convergence, leading to the suggestion that temporal synchrony binds cortical areas (15). This hypothesis was first supported by phase-locked unit firing and local field potentials (LFP) at 40-60Hz evoked by simple visual stimuli in the anesthetized cat primary visual cortex at distances <7mm (16). Although some further studies found similar results in other cortical areas, behaviors and species, as would be expected under the binding-by-synchrony hypothesis (17, 18), others have been less successful (19). Synchronous high gamma oscillations have also been criticized as providing no mechanism for neuronal interaction beyond generic activation (19, 20).

Here, using human intracranial stereoelectroencephalography (SEEG) recordings, we find that ripples co-occur, and remarkably, phase-synchronize across all lobes and between both hemispheres, with little decrement, even at long distances. Furthermore, ripple co-occurrence is enhanced between cortical sites as well as between the cortex and hippocampus preceding successful delayed recall. Co-rippling was progressively above that expected as it involved a larger proportion of sites, and this led to progressively stronger phase-locking. Single-unit firing increased during, and phase-locked to, cortical ripples, providing a basic requirement for ripples to enhance communication via gain-modulation and coincidence detection. Enhanced communication was supported by our finding of increased high-frequency correlation between even distant co-rippling regions. These characteristics suggest that distributed, phase-locked cortical ripples possess the properties that may allow them to help integrate different components of a memory. More generally, ripples may help to ‘bind’ different aspects of a mental event encoded in widespread cortical areas into a coherent representation.

## Results

### Ripple detection during NREM and waking

Ripples were detected using intracranial cortical and hippocampal recordings in 17 patients (STable 1) undergoing monitoring to localize seizure foci during NREM and waking. Bipolar contact derivations were used to measure LFPs. Ripples were detected only on channels in non-lesional, non-epileptogenic areas. Ripples were required to have 3 or more cycles of increased peak 70-100Hz analytic amplitude that did not contain epileptiform activity or artifacts. Ripples were also required not to occur at the time of putative interictal spikes detected on any other channel to exclude events that could be spuriously coupled due to epileptiform activity.

Cortical ripples were detected in all lobes of both hemispheres during NREM and waking (Fig.1A-B, *N*=273 channels). Cortical and hippocampal ripples were consistently ~70-85ms long ~90Hz oscillations. Specifically, cortical ripples during NREM had an average and standard deviation (across channel averages) frequency of 89.1±0.8Hz, density of 8.4±2.7min^-1^, amplitude of 5.1±2.4μV, and duration of 76.5±10.1ms. During waking, cortical ripples had a frequency of 89.5±0.7Hz, density of 5.8±3.4min^-1^, amplitude of 7.2±3.9μV, and duration of 66.6±7.2ms (Fig.1C-D; *N*=273 channels). Hippocampal ripples during NREM had an average and standard deviation frequency of 85.7±2.0Hz, density of 7.0±4.8min^-1^, amplitude of 17.4±7.8μV, and duration of 87.5±9.0ms, and during waking had a frequency of 88.2±1.4Hz density of 7.4±5.5min^-1^, amplitude of 18.0±9.6μV, and duration of 69.7±7.9ms (*N*=28 channels). A standard deviation of <1Hz in ripple frequency in all circumstances suggests the possibility that local channel and network properties underlie a consistent resonant frequency.

**Fig. 1.**
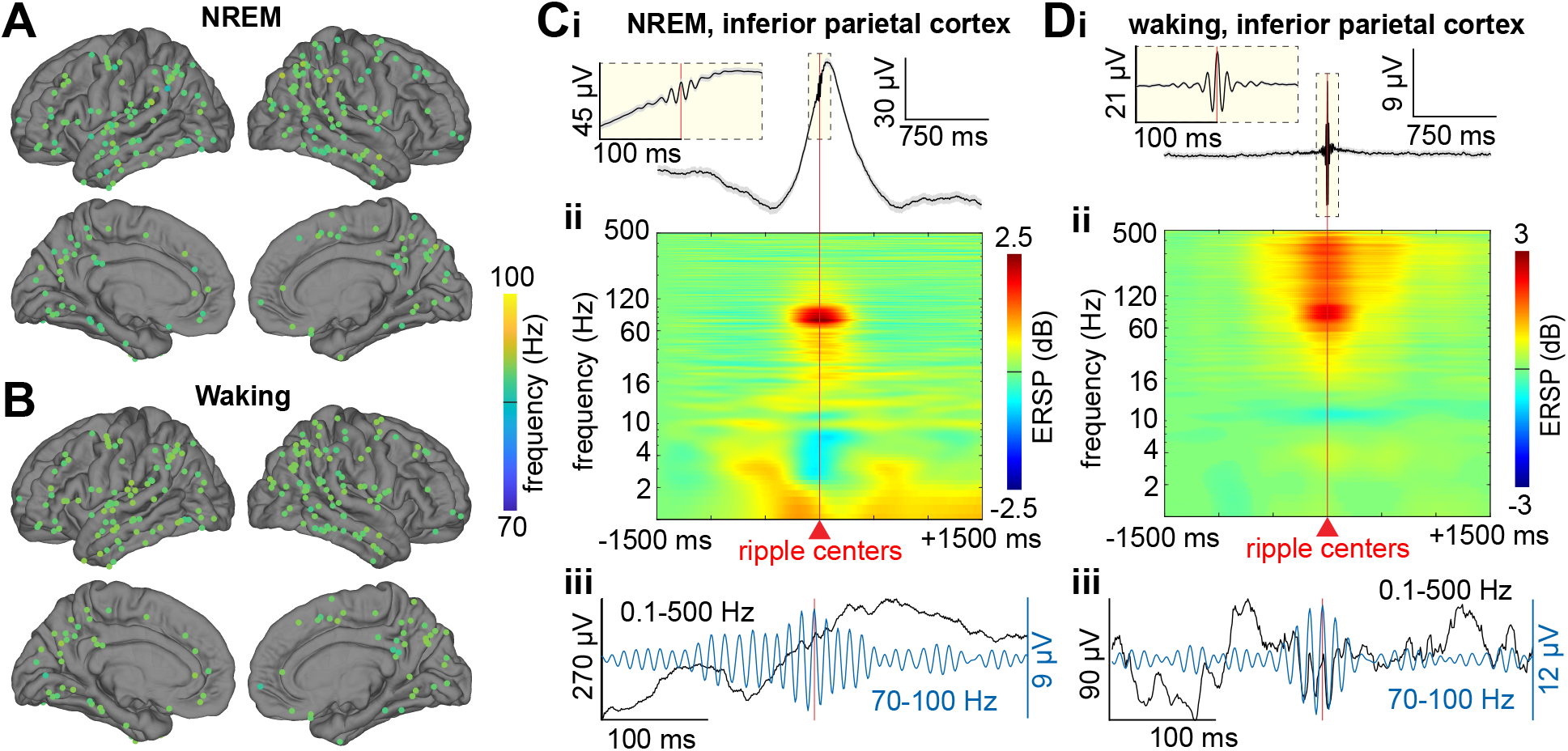
Cortical ripple generation during NREM and waking. (**A**) Cortical ripple oscillation frequencies across the cortex during NREM. Each marker depicts the average ripple frequency in a given channel (*N*=273 bipolar channels from SEEG patients S1-17). Color range spans the ripple band used for detection (70-100Hz). Note the highly consistent frequencies across lobes and between hemispheres. (**B**) Same as (A) except for waking. Note the highly consistent frequencies between NREM and waking. (**C**) Average broadband LFP (**i**) and time-frequency (**ii**) across cortical ripples, and example broadband ripple (**iii**) unfiltered single sweep (black), and 70-100Hz bandpass (blue) during NREM, in inferior parietal cortex. (**D**) Same as (A) except during waking. Error shows SEM. ERSP=event-related spectral power, NREM=non-rapid eye movement sleep, SEEG=stereoelectroencephalgography.

### Cortical ripples couple and co-occur across widespread regions

We hypothesized that ripples couple and especially co-occur between cortical sites, which would be crucial for widespread information integration. We discovered that cortical ripples frequently and strongly couple (occur within ±500ms of each other) during both NREM (*N*=4487/4550 significant channel pairs, randomization test, post-false discovery rate (FDR) *p*<0.05) and waking (*N*=4478/4550), between all cortical areas sampled (Fig.2A, Table 1), including between hemispheres. The proportion of cortico-cortical channel pairs that were significantly coupled during NREM (98.6%) was not significantly different than that during waking (98.4%; *p*=0.44, χ^2^=0.61, df=1). See SFig.2A-B for individual patients.

**Fig. 2.**
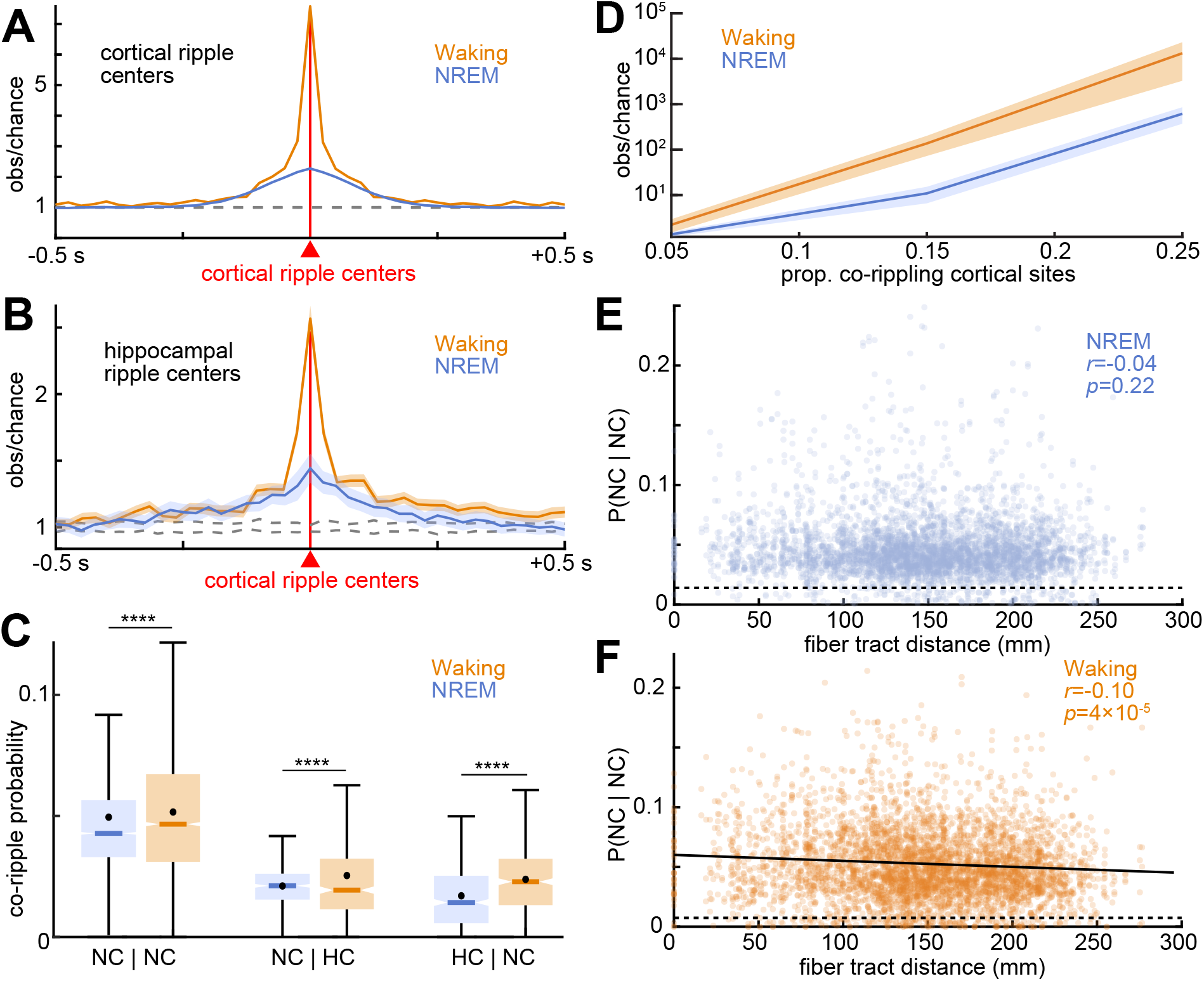
Cortico-cortical and hippocampo-cortical ripples couple and co-occur during waking and NREM. (**A**) Cross correlograms of ripples between all possible cortical sites reveals strong ripple coupling (occurring within 500ms of each other without necessarily overlapping) between nearly all sites during NREM (*N*=4487/4550 significant channel pairs) and waking (*N*=4478/4550; post-FDR *p*<0.05, randomization test). Dashed lines show 99% confidence interval of the null distribution (200 shuffles/channel pair). (**B**) Same as (A) except crosscorrelogram of hippocampal ripples relative to cortical ripples (NREM: *N*=133/461; waking: *N*=401/461). (**C**) Conditional co-occurrence probabilities of cortico-cortical and hippocampo-cortical ripples overlapping for at least 25ms (i.e., probability of a ripple in a particular channel given a ripple in another particular channel) are greater during waking than NREM (****post-FDR *p*<0.0001, two-sided paired *t*-test). (**D**) Observed over chance cortical ripple co-occurrence (logarithmic scale) increases with the number of sites co-rippling. (**E**) Cortical ripple co-occurrence probabilities during NREM are stable with increasing intervening fiber tract distance (linear mixed-effects with patient as random effect). Dashed line indicates chance. When two channels were in the same parcel the fiber tract distance was defined as 0. (**F**) Same as (E) except for waking. Fit is linear least-squares regression. Error shows SEM. FDR=false discovery rate.

**Table 1.** Cortical ripple coupling with cortical or hippocampal ripples: frequency and order. <u>*Significant Modulation:*</u> Proportion of channel pairs with a significant increase in the conditional probability of a ripple occurring in the first channel given that one occurred within ±500ms in the second (e.g., Hipp-R | Cort-R refers to hippocampal ripples occurring within ±500ms relative to cortical ripples at *t*=0; one-sided randomization test, 200 shuffles, 25ms nonoverlapping bins, 3 consecutive bins each with post-FDR *p*<0.05 required for significance). During NREM, conditional probabilities are shown separately for those hippocampal ripples associated with sharpwaves (Hipp-SWR) and sleep spindles (Hipp-SSR) as well as all ripples (Hipp-R). <u>*Significant Sidedness:*</u> Those with significant modulations that had significant sidedness preference around *t*=0 (post-FDR *p*<0.05, two-sided binomial test, expected=0.5, −500 to −1ms vs. 1 to 500ms). <u>*Cortical Ripple Leading:*</u> Those with significant sidedness around 0 that had cortical ripples leading (according to counts within −500 to −1ms vs. 1 to 500ms). During NREM, Hipp-R and Hipp-SWR led Cort-R, and during waking, Cort-R led Hipp-R (*=*p*<0.05, two-sided binomial test, expected value=0.5), resulting in a significant preference for cortical ripples to lead hippocampal ripples in waking vs. NREM (*p*<0.00001, χ^2^=51.59, df=1). See STable 3 for results from individual patients.

| Ripple \| Ripple | Significant Modulation | Significant Sidedness | Cortical Ripple Leading |
| --- | --- | --- | --- |
|  | NREM |  |  |
| Cort-R \| Cort-R | 98.62% (4487/4550) | 3.50% (157/4487) | N/A |
| Hipp-R \| Cort-R | 28.85% (133/461) | 31.58% (42/133) | 33.33% (14/42)* |
| Cort-R \| Hipp-R | 29.28% (135/461) | 31.11% (42/135) | 33.33% (14/42)* |
| Hipp-SWR \| Cort-R | 19.74% (91/461) | 45.05% (41/91) | 24.39% (10/41)* |
| Cort-R \| Hipp-SWR | 20.61% (95/461) | 44.21% (42/95) | 21.43% (9/42)* |
| Hipp-SSR \| Cort-R | 12.15% (56/461) | 17.85% (10/56) | 50% (5/10) |
| Cort-R \| Hipp-SSR | 11.71% (54/461) | 22.22% (11/54) | 41.67% (5/12) |
|  | Waking |  |  |
| Cort-R \| Cort-R | 98.42% (4478/4550) | 8.13% (364/4478) | N/A |
| Hipp-R \| Cort-R | 86.98% (401/461) | 22.44% (90/401) | 84.44% (76/90)* |
| Cort-R \| Hipp-R | 87.42% (403/461) | 22.33% (90/403) | 84.44% (76/90)* |

Critically, we found that short latency coupling led to co-occurrence (≥25ms overlap) of these ~70ms long ripples, with slightly but significantly higher probabilities during waking than NREM (Fig.2C). The percent of cortico-cortical channel pairs with significant ripple co-occurrence was very high and nearly equal during NREM (*N*=2218/2275, 97.5%) vs. waking (*N*=2225/2275, 97.8%). We also found that the co-occurrence probabilities of cortico-cortical ripples across channels were correlated between NREM and waking (SFig.3A; *r*=0.41, *p*=7×10^−180^, significance of *r*). In sum, nearly all cortico-cortical channel pairs couple and co-occur above chance during both NREM and waking.

### Hippocampal and cortical ripples couple and co-occur, especially during waking, but at lower rates than cortico-cortical

As previously found (8), ripples in many hippocampo-cortical channel pairs were also significantly coupled (Fig.2B, Table 1) during both NREM (*N*=133/461) and waking (*N*=401/461), but at a significantly lower rate than cortico-cortical pairs in both states (NREM: *p*<0.00001, χ^2^=2832.0, df=1; Waking: *p*<0.00001, χ^2^=213.3, df=1). Unlike cortico-cortical pairs, the proportion of significant hippocampo-cortical pairs was higher during waking (87.0%) than NREM (28.9%, *p*<0.00001, χ^2^=319.6, df=1). See SFig.2C-D for individual patients. These findings of cortico-cortical and hippocampo-cortical couplings were maintained when only cortical and hippocampal channels that were free of interictal spikes at any time were analyzed (STable 2).

Overlapping co-occurrence of human ripples between cortex and hippocampus do not appear to have been studied *per se,* but would also be expected to occur given their coupling. Indeed, we found significant co-occurrence of ripples in 81.1% (*N*=374/461) of hippocampo-cortical site pairs during waking, and 34.7% (*N*=160/461) during NREM. Higher hippocampo-cortical cooccurrence during waking compared to NREM was significant (*p*<0.00001, χ^2^=203.8, df=1). In addition, these percentages for hippocampo-cortical co-occurrences (81.1% waking; 34.7% NREM) are significantly lower than for cortico-cortical site pairs (97.5% waking; 97.8% NREM) (NREM: *p*<0.00001, χ^2^=1328.8, df=1; waking: *p*<0.00001, χ^2^=1250.1, df=1). Thus, ripples in hippocampal and cortical sites do couple and co-occur, but at a substantially lower rate than between cortical sites.

### Cortical ripples lead hippocampal ripples during waking

Some memory models posit that information is transferred during waking from cortex to hippocampus for memory encoding. During waking, 22.4% (90/401) of channel pairs had a significant order preference (Table 1; post-FDR *p*<0.05, two-sided binomial test, expected value=0.05). Among these significant pairs, cortical ripples led hippocampal ripples in 84.4% (76/90). During NREM, 31.6% (42/133) of the hippocampo-cortical pairs that were significantly coupled had a significant sidedness preference, where one channel’s ripples led the other’s. While hippocampal ripples led cortical ripples in 66.7% (28/42) of these significant pairs during NREM, this effect was largely driven by one patient (STable 3), unlike the effect of cortical ripples leading hippocampal ripples during waking, which was consistent across the majority of patients (STable 3). The overall preference for cortical ripples to lead hippocampal ripples during waking compared to NREM was highly significant (*p*=4.3×10^-9^, χ^2^=34.5, df=1).

### Hippocampal sharpwave ripples lead cortical ripples during NREM

While the results above show that cortical ripples precede hippocampal ripples during waking, their order during NREM appears to be a mixed picture, which we hypothesized depends on whether the hippocampal ripples occur in the context of a sharpwave or spindle (2, 21). We found that hippocampal sharpwave-ripples significantly preceded cortical ripples by ~250ms (SFig.1A; *p*=0.002, two-sided binomial test, expected value=0.5), whereas spindle-ripples were concurrent (SFig.1B; *p*=1). These results reinforce a previous suggestion that sharpwave- and spindle-ripples make sequential contributions to consolidation. Overall, as predicted by models of hippocampo-cortical interaction for memory, hippocampal ripples usually lead cortical during sleep, and cortical usually lead hippocampal during waking.

### Cortical ripple co-occurrence is facilitated through activation across multiple sites

In each patient, we found that when two cortical sites co-rippled, one or more additional sites may join them (SFig.5). To test if two cortical sites co-rippling made it more likely for other sites also to co-ripple, we computed a χ^2^ test of proportions for all possible groups of three cortical channels under the null hypothesis that the co-occurrence of channels A and B has no relation to the co-occurrence of A and C. We found significantly increased co-occurrence of the third site in an average of 14.1% of triplets during NREM and 38.8% during waking (patient specific results reported in STable 4, χ^2^ test of proportions, FDR-corrected *p*-values across channel-triplets within patients).

In further support that co-rippling is facilitated through activation across multiple sites, we found that the number of ripple co-occurrences relative to chance increased with the number of locations co-rippling (>2) (Fig.2D). The increase was pronounced, such that the observed number of co-occurrences relative to baseline of 25% of all channels was increased by a factor of ~10^4^ during waking and ~5×10^3^ during NREM. Thus, co-rippling appears to beget more co-rippling, suggesting the possibility of self-reinforcing spread.

### Cortical ripples co-occur robustly across distance

Binding by ripples requires that they co-occur across widespread cortical areas. We compared conditional probabilities of cortico-cortical ripple co-occurrences (i.e., the probability of a ripple in one cortical site given a ripple in another cortical site, requiring ≥25ms overlap) against whitematter streamline distances between cortical sites. Streamline distances were computed using the 360 parcels of the HCP-MMP1.0 atlas (22), as determined by probabilistic diffusion MRI tractography (23), and are population averages (24). We found that cortico-cortical co-rippling probability did not decrease with fiber tract distance during NREM (Fig.2E, *r*=-0.04, *p*=0.22, linear mixed-effects with patient as random effect), but rather was maintained up to 25cm separation, across lobes and between hemispheres. During waking, co-rippling probabilities were also maintained across this distance range, albeit with a weak but significant negative linear relationship (Fig.2F, *r*=-0.10, *p*=4×10^-5^). See SFig.4A-B for individual patients.

### Ripples in all cortical areas co-occur with hippocampal ripples

It was previously reported that ripple coupling occurs between the parahippocampal gyrus and 16.4% of lateral temporal electrodes but only 3.3% of Rolandic (8), perhaps reflecting the anatomical location of the hippocampus at the apex of the cortical hierararchy (25). However, we found that hippocampal ripples co-occurred with ripples in all cortical areas at approximately equal rates in both NREM and waking. Taking the myelination index as a measure of position in the cortical hierarchy (association areas are less myelinated (26)), we found a weak but significant effect during NREM but not waking. During NREM, hippocampo-cortical cooccurrence was positively correlated with myelination (SFig.6; *r*=0.15, *p*=0.01), indicating that hippocampal ripples are slightly more likely to co-occur with ripples in primary cortical areas.

### Cortico-cortical and hippocampo-cortical ripple co-occurrence precedes recall

If ripples in different cortical regions bind the elements of a memory, it would be expected that cortical ripples co-occur preceding cued recall, which requires that those elements be coactivated. To test this hypothesis, we analyzed paired-associates memory task data from 5 SEEG patients (Fig.3A, STables 1,5). Preceding delayed cued recall, there was a significant increase in cortical ripple occurrence (*p*=1×10^-11^; linear mixed-effects models with patient as random effect) of 330% of chance (computed on a trial-wise basis, thus chance was determined separately for immediate and delayed recall), and an even greater increase in cortico-cortical ripple co-occurrence of 758% compared to chance (*p*=0.0004; Fig.3B-E). Furthermore, cortical ripple occurrence (*p*=0.002) and cortico-cortical (*p*=0.002) and hippocampo-cortical (*p*=0.008) ripple co-occurrence modulations were greater preceding delayed vs. immediate recall, which shared the same stimuli and responses. Finally, cortico-cortical (*p*=0.04) and hippocampo-cortical (*p*=0.004) co-rippling was enhanced preceding correct vs. incorrect delayed recall, which was not the case for cortical (*p*=0.08) or hippocampal (*p*=0.94) ripples generally. Notably, delayed but not immediate recall of paired associates is severely impaired by hippocampal damage (12). These data demonstrate increased co-rippling between cortical sites and between the hippocampus and cortex during hippocampal-dependent retrieval of novel combinations of previously unrelated items. This finding supports the hypothesis that hippocampo-cortical and cortico-cortical co-rippling may contribute to the reconstruction of declarative memories in humans.

**Fig. 3.**
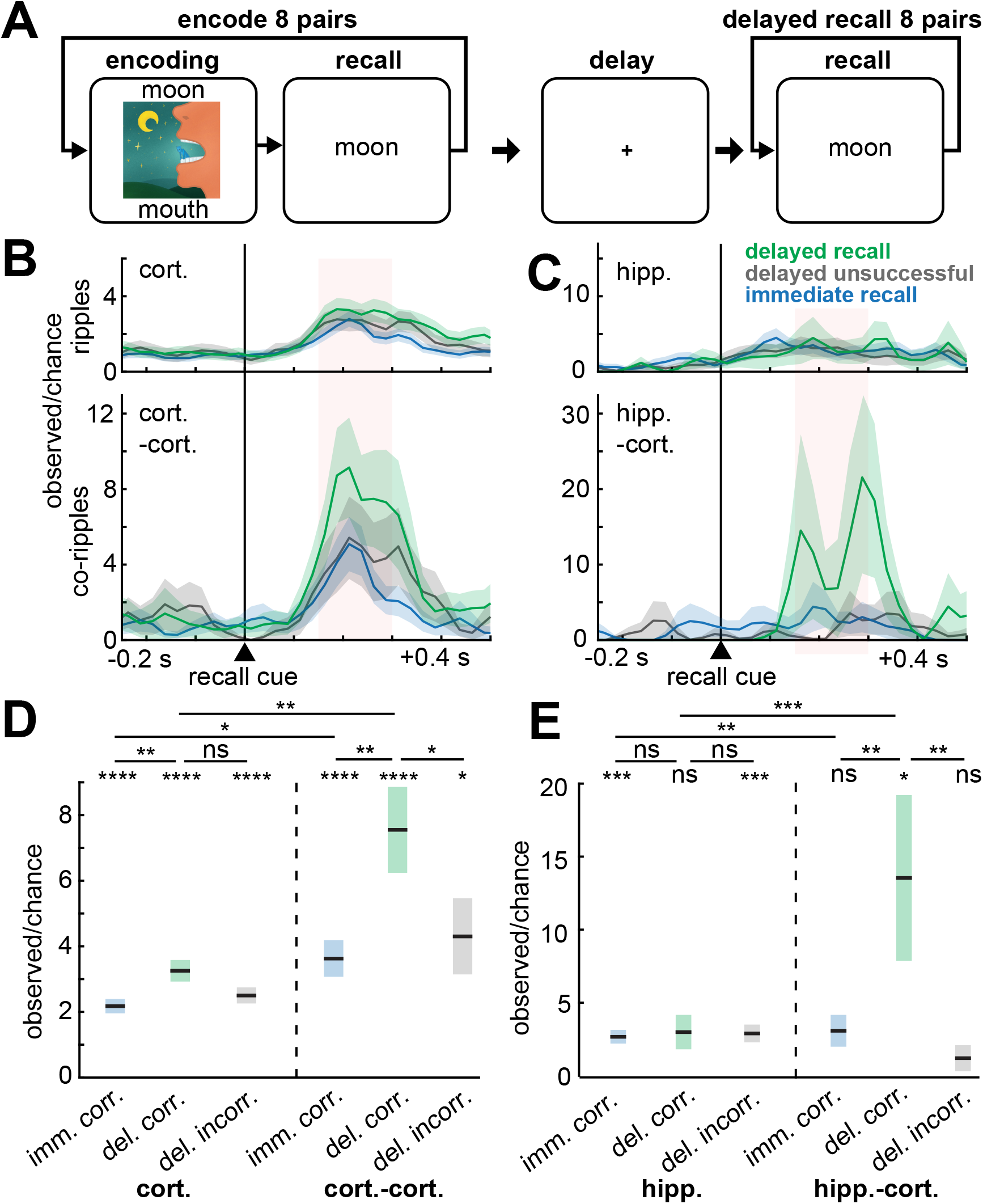
Cortico-cortical and hippocampo-cortical ripple co-occurrences increase preceding recall. (**A**) Schematic of paired-associates memory task. Patients learned word pair associations and were subsequently cued with the first word to recall the second immediately following learning and then after a ~60s delay with distraction. (**B**) Cortical and cortico-cortical co-rippling increases following the stimulus cue triggering recall (delayed *N*=365 trials, immediate *N*=698 trials, patients S18-22). (**C**) Same as (B) except for hippocampal ripples and hippocampo-cortical co-ripples (delayed *N*=90 trials, immediate *N*=304 trials, patients S19,22). **(D-E**) Quantification of (B-C) during the pink shaded 150-300ms interval following stimulus cue onset preceding correct recall in the immediate or delayed condition, or incorrect recall (no attempt or incorrect response) in the delayed condition. Note that cortico-cortical and hippocampo-cortical co-occurrences preceding correct delayed recall have the greatest increases. Errors show SEM. Post-FDR \**p*<0.05, \*\**p*<0.01, \*\*\**p*<0.001, \*\*\*\**p*<0.0001, linear mixed-effects models with patient as random effect.

Since brain state could affect recall performance, we tested whether there was a difference in alpha (7-13 Hz) analytic amplitude between correct and incorrect responses. We found no significant difference in the mean alpha analytic amplitude across cortical channels within a ±1 s window around response times for correct vs. incorrect for any of the 5 patients with paired-associates task data (*p*=0.24-0.64 and *t*=0.36-1.6, one-sided two-sample *t*-test, FDR-corrected *p*-values for multiple patients).

### Cortical ripples phase-lock across widespread regions

Phase-locked oscillations in widespread locations have been hypothesized to underlie integration (‘binding’) of different components of events across the cortical surface. For each channel pair, we calculated the phase-locking value (PLV), a measure of the consistency of 70-100Hz phases between sites, independent of amplitude (27), across all of their co-ripple events in either NREM or waking. Please note that consistency is measured between different co-ripples, at each 1ms bin relative to co-ripple center, not within co-ripples. We found significant PLV of co-occurring ripples between all sampled cortical regions, including between hemispheres, in both states (Fig.4A-C). Channel pairs with significant PLV were more frequent during NREM than waking (Fig.4C, STable 6; post-FDR *p*<0.05, randomization test; nonsignificant results in SFig.7A-B). An example of consistent phase between two cortical locations across two co-ripples is shown in Fig.4A. For each channel pair, PLV was measured at each latency relative to the center of their co-ripple (Fig.4B), and these time-courses were averaged across all significant channel pairs (Fig.4C). The increased PLV lasted for the entire period when the sites were co-rippling, and arose abruptly from baseline. PLV time-course was remarkably similar across channel pairs. During sleep 2106/2275 cortico-cortical channel pairs had more than 40 co-occurring ripples, required for a reliable PLV estimate. Of these, 26.3% (554/2106) had significant PLV modulations. During waking, 1939/2275 had more than 40 cooccurring ripples, and 13.9% (269/1939) of these had significant PLV modulations (STable 6). Like what we found for co-occurrences (SFig.3A), cortico-cortical co-ripple peak PLVs across channel pairs were positively correlated between NREM and waking (SFig.3B; *r*=0.20, *p*=4×10^−22^, significance of *r)*. In summary, distant pairs of sites throughout the cortex were found to have consistent phases during co-ripples in both NREM and waking.

**Fig. 4.**
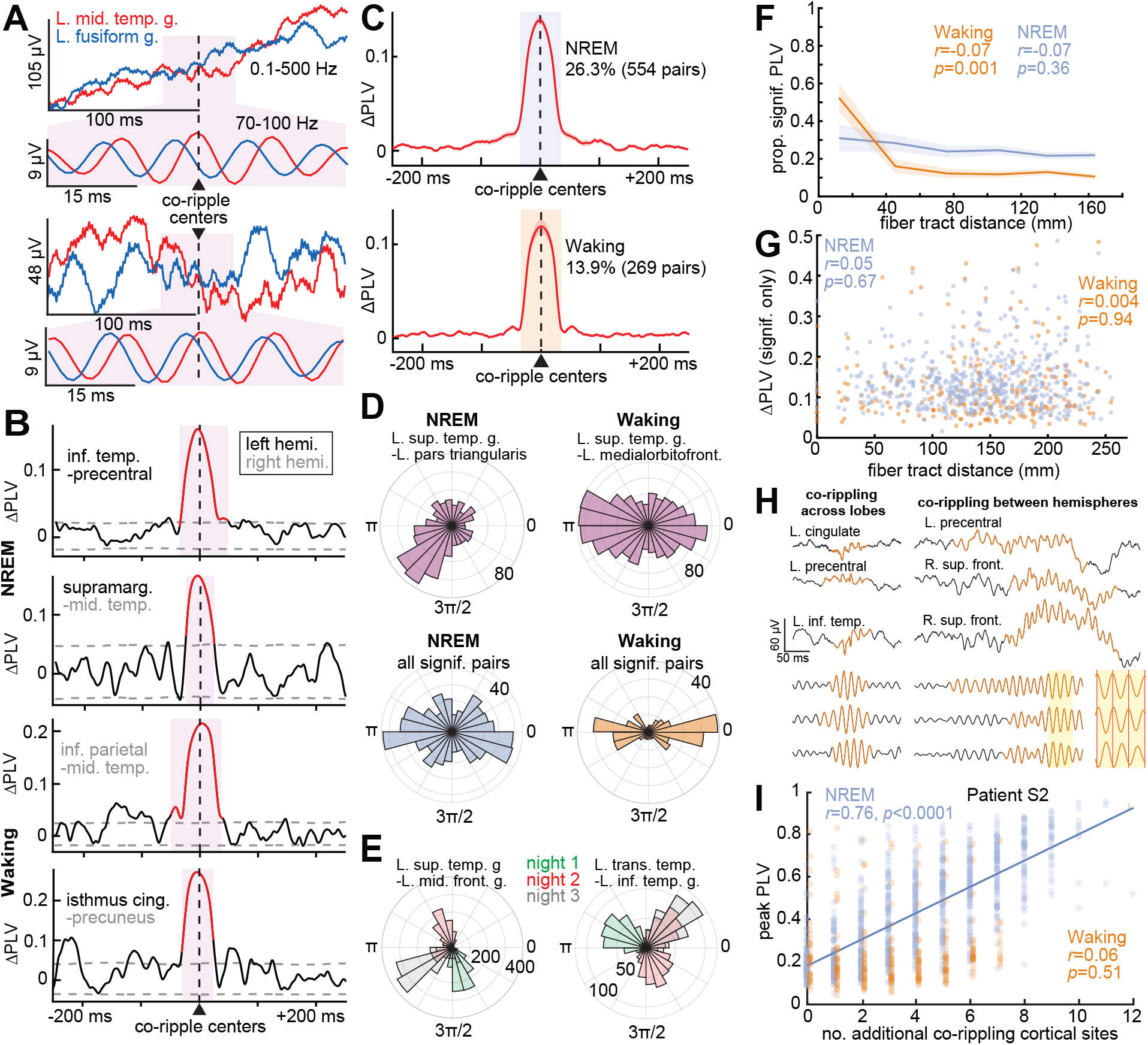
Ripples phase-lock across wide separations in the cortex. (**A**) Two example pairs of co-occurring ripples in broadband LFP and 70-100Hz bandpass. A consistent phase-lag from the left middle temporal gyrus (red) to left fusiform gyrus (blue) is evident in the expanded bandpassed (70-100Hz) 50ms long traces centered on the co-ripple (pink background). Similar phase-lags were also present across other co-ripples between these sites, resulting in a significant PLV. (**B**) Example 70-100Hz ΔPLV time-courses calculated between ripples cooccurring between ipsilateral and contralateral cortical sites (≥25ms overlap) in NREM and waking. Red shows significant modulation (post-FDR *p*<0.05, randomization test, 200 shuffles/channel pair). (**C**) Average ΔPLVs (relative to −500 to −250ms) for cortical channel pairs with significant PLV modulations. A greater percentage of pairs had significant co-ripple PLV modulations during NREM (26.3%) than waking (13.9%). (**D**) Polar histograms of waking and NREM 70-100Hz phase-lags across co-ripples for two example cortico-cortical pairs with significant PLV modulations (top) and across channel pair circular means (bottom). The magnitude of each pie-wedge corresponds to the number of co-ripples (top) or number of channel pairs (bottom) with the indicated phase-lag. Cortico-cortical phase-lags had a significant preference for ~0 or ~π during waking compared to NREM based on the counts within 0±π/6 or π±π/6 vs. outside these ranges for NREM vs. waking (*p*=5×10^-8^, χ^2^=29.8, df=1). (**E**) Example co-ripple phase-lag distributions for different sleep nights. The dominant phase-lag for co-ripples changes across nights (color-coded). (**F**) Proportion of channel pairs with significant PLVs has a weak, non-significant decrease over distance during NREM, but a significant decrease, notably at shorter distances, during waking (linear mixed-effects with patient as random effect). See SFig.4C-D for individual patients. (**G**) ΔPLV does not decrement with intervening fiber tract distance for channel pairs with significant co-ripple PLV modulations (linear mixed-effects with patient as random effect). See SFig.4E-F for results from individual patients. (**H**) Single sweep broadband LFP and normalized 70-100Hz bandpass show waking ripples (orange) co-occurring across lobes (left) and between hemispheres (right). Note the consistency of ~0 or ~π phases in the shaded inset. (**I**) Peak PLV correlates with the number of additional cortical sites co-rippling during NREM in a sample patient (*N*=10/17 patients significant, post-FDR *p*<0.05, significance of the correlation coefficient) more often than waking (*N*=3/17). Fit is linear least-squares regression. PLV=phase-locking value. The PLVs between the indicated channels measure the consistency of phase between the those channels at each latency relative to ripple peak across all instances of ripples co-occurring between those channels in NREM or waking.

**Fig. 5.**
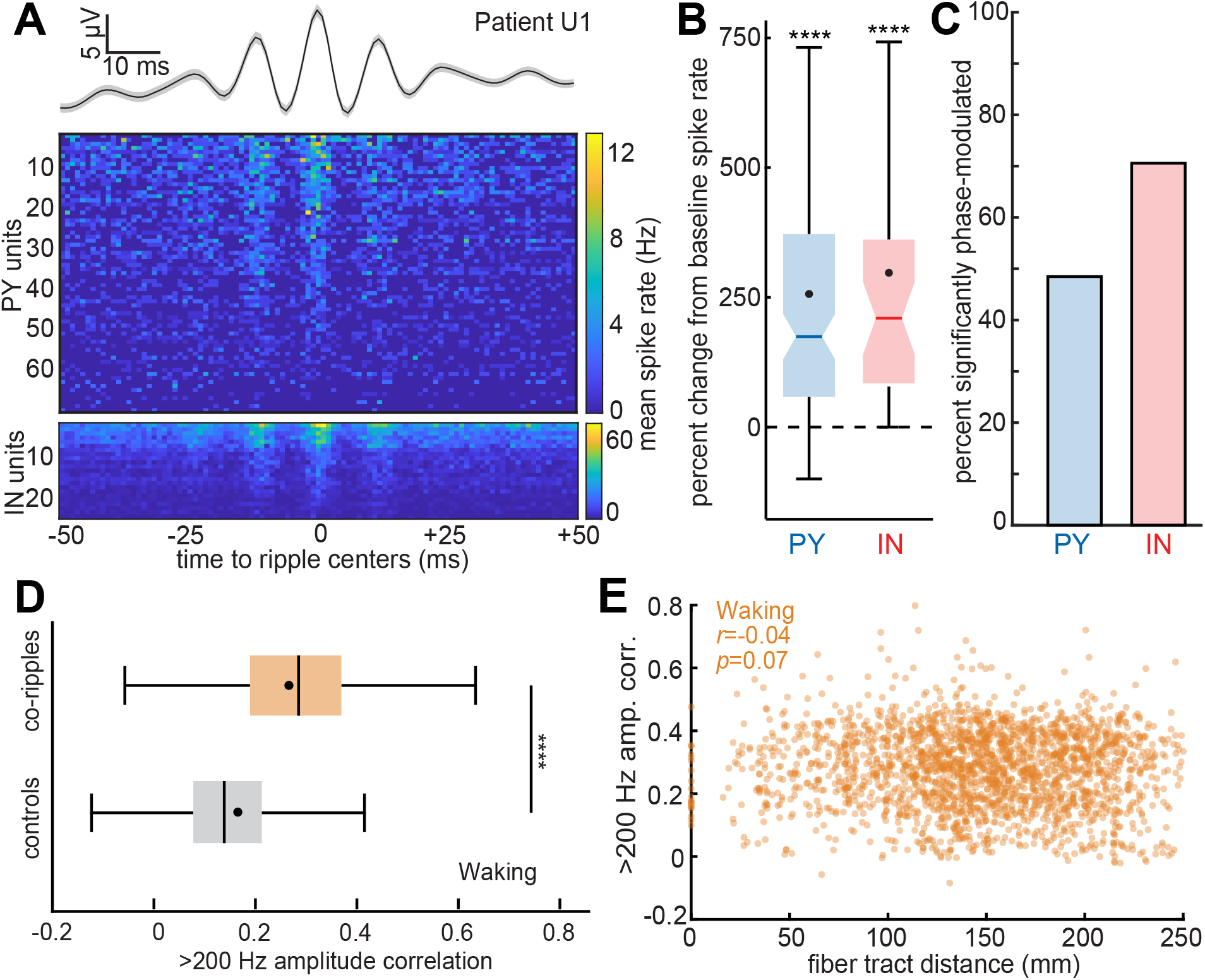
Cortical ripples modulate local single unit spiking and synchronize high-frequency activity between distant regions. (**A**) Average cortical ripple broadband LFP and associated raster plot of pyramidal (PY) and interneuron (IN) mean spike rates across ripples during NREM. Note the phase-coupling of both PY and IN spiking to the ripple peaks. (**B**) Single unit spike rates increase during cortical ripples compared to baseline, which was comprised of randomly selected epochs in between ripples matched in number and duration (PY: *N*=127, mean=255%; IN: *N*=38, mean=297%; patients U1-3). (**C**) Single units are significantly phase-modulated by cortical ripples (PY: *N*=32/66 significant; IN: *N*=24/34 significant; binomial test between phases within 0±π/2 vs. π±π/2, expected value=0.5, minimum 30 spikes per unit across ripples). (**D**) Correlation of the >200Hz analytic amplitude, a proxy for unit spiking, increases during waking co-ripples compared to randomly selected preceding control periods (within −10 to −2 s) matched in number and duration (*N*=2275 SEEG channel pairs). ****p<0.0001, two-sided paired *t*-test. (**E**) Average >200Hz amplitude correlation between co-ripples for each cortical channel pair does not decrement with fiber tract distance.

### Phase-locked co-ripples have a broader range of phase-lags in NREM compared to waking and may vary across nights

Having demonstrated significant phase-locking between cortical co-rippling sites, we evaluated the distribution of the circular mean phase angles across site pairs. We found that during NREM the average phase angles for different cortical site pairs that had significant co-ripple phaselocking had a fairly even distribution from 0 to 2π radians (Fig.4D bottom left). However, during waking, co-ripple phase-lags across pairs tended to be ~0 or ~π (Fig.4D bottom right). This difference was significant (*p*=5×10^-8^, χ^2^=29.8, df=1; using counts within 0±π/6 or π±π/6 vs. outside these ranges for NREM vs. waking). This observation suggests a greater tendency toward zero phase-lag during waking (we provide evidence below that the ~0 and ~π lags are functionally equivalent and may be due to a variability in fine-scale electrode contact placement relative to cortical layers). This may be related to the greater tendency of ripples to couple at near zero latency during waking (Fig.2A).

We also tested whether phase-lags during NREM varied across nights. For each patient with multiple sleep nights (*N*=16), we compared the co-ripple phase-lags between all possible pairs of sleep nights within channel pairs that had significant PLV modulations. We found that 51.8% (1256/2426) of such pairs were significantly different in their phase-lags between nights (Fig.4E; post-FDR *p*<0.05, Watson-Williams test; minimum 30 co-ripples per night per channel pair). These differences in co-ripple phase-lags between particular cortical sites suggest that they may participate in different networks across nights, as may happen, for example, when re-activating cortical representations associated with different memories. Similar phenomena have been noted in large-scale cortical models (28).

### Ripples phase-lock robustly across long distances

Since declarative memories characteristically unite disparate elements that are encoded in widespread cortical locations in both hemispheres, any neurophysiological process supporting their encoding, consolidation, or retrieval must likewise function across those distances. Thus, given that cortico-cortical ripple co-occurrences do not decrement with distance (tested up to 200mm, Fig.2E-F), we hypothesized that cortico-cortical co-ripple phase-locking also does not decrement with distance. Indeed, we found that, like ripple co-occurrences, the proportion of channel pairs with significant PLV modulations (Fig.4F; *r*=-0.07, *p*=0.36, linear mixed-effects with patient as random effect) and the magnitude of these PLV modulations (Fig.4G; *r*=0.05, *p*=0.67) did not significantly decrement with intervening fiber tract distance during NREM. During waking, there was a significant decrement in the proportion of channel pairs with significant PLV modulations with distance, especially at shorter distances (Fig.4F; *r*=-0.07, *p*=0.001), but no decrement in the magnitude of significant channel pair PLV modulations with distance (Fig.4G; *r*=0.004, *p*=0.94). Thus, the physiological action supported by co-rippling may span the entire cortical surface.

### Phase-locking increases with more cortical sites co-rippling

Ripples often phase-locked across multiple sites, including between hemispheres (Fig.4H), and as described above we found that two sites co-rippling made it more likely that a third site was co-rippling. These findings raise the question of whether activation of widespread co-rippling networks facilitates greater network synchronization. We found a strong positive linear correlation between the number of sites co-rippling and the cortico-cortical co-ripple phaselocking peak amplitude (Fig.4I). These results were again found when the ΔPLV (peak PLV minus baseline PLV) was used instead of the peak PLV (ΔPLV: *N*=7/17 patients significant during NREM, *N*=1/17 significant during waking, post-FDR *p*<0.05, significance of the correlation coefficient). To test whether more co-rippling was simple due to greater ripple amplitude, we measured the average 70-100 Hz analytic amplitude of the co-ripples, and found that only 2/17 patients in NREM and 0/17 patients in waking had significant correlations, (post-FDR p<0.05, significance of the correlation coefficient), and the significant correlations were both negative, demonstrating that more co-rippling was not due to greater amplitude. In sum, the co-occurrence of ripples promotes further co-occurrence, which enhances phase-locking.

### Cortico-cortical co-ripples have consistent phase-lags across successive cycles

We hypothesized that a site pair would be effectively ‘phase-locked’ across successive co-ripple cycles because the ripple oscillation frequency of ~90Hz is so similar across ripples and locations. For each co-ripple from each cortico-cortical channel pair we computed the PLV across the lags between the two ripples using their 5 peaks closest to the co-ripple time center, and found significant within-ripple phase-locking for 98.3% of the 807,213 co-ripples during NREM and 98.0% of the 1,348,696 co-ripples during waking (*p*<0.05; randomization test, *N*=1000 random phase-lags, FDR-correction across co-ripples). Thus, within-ripple phaselocking is present in nearly all cortico-cortical co-ripples.

### Hippocampo-cortical pairs rarely if ever phase-lock

Cortico-cortical phase-locking could be driven through a network of coupled cortical oscillators, a central driving mechanism, or a combination (Fig.6A). Since hippocampal ripples strongly cooccur with cortical ripples (Fig.2B), we investigated whether the hippocampus drives phaselocking of ripples in the cortex by testing if there is phase-locking between hippocampo-cortical co-ripples. For hippocampo-cortical pairs during sleep, 277/461 had more than 40 co-occurring ripples, and 1.4% (4/277) of these had significant PLV modulations. During waking, 333/461 of the hippocampo-cortical channel pairs had more than 40 co-occurring ripples, and 0.3% (1/333) of these had a significant PLV modulation (SFig.7C-F; STable 6). Examination of electrode trajectories suggested that the hippocampal contacts with significant PLV with cortical sites were probably located in the subicular complex rather than the hippocampus proper, consistent with the finding in mice that ripple propagation from hippocampus to retrosplenial cortex is via the subiculum (10). Thus, it is unlikely that cortico-cortical ripple phase-locking is driven by common inputs from a hippocampal ripple, which supports the hypothesis that cortico-cortical co-ripple phase-locking is driven intracortically.

**Fig. 6.**
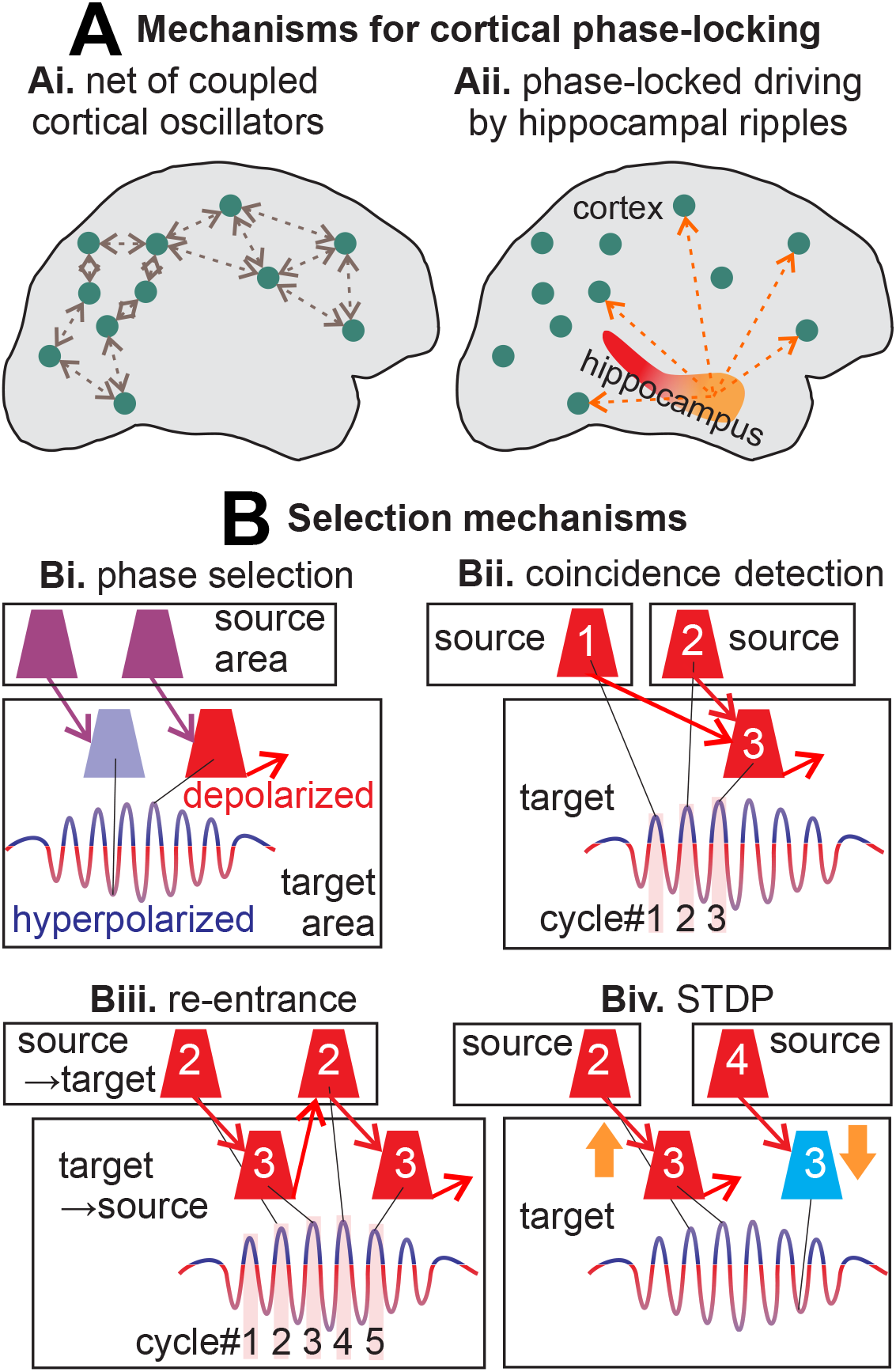
Mechanisms of phase-locked networks of cortical ripples and their selection. (**A**) Potential mechanisms for ripple phase-locking across broad cortical regions: (**Ai**) multiple cortico-cortical re-entrant connections between the cortical oscillators; (**Aii**) distributed driving oscillations from a subcortical location (e.g., hippocampal ripples). (**B**) Multiple synergistic mechanisms enabled by cortico-cortical co-rippling could select a spatiotemporal neural network. (**Bi**) Spikes arriving at the hyperpolarized phase will be ineffective relative to those arriving at the depolarized phase in eliciting spikes in the rippling target area. (**Bii**) Coincident spikes arriving from two source areas that are phase-locked with the target area are more likely to trigger an action potential, especially if they arrive in the depolarized phase. Note that conduction times between areas does not need to match the phase-lag but could equal the phase-lag plus multiples of the cycle time (e.g., cell #1→3). (**Biii**) Re-entrant activation across cycles could also result from reciprocal connections. (**Biv**) The synapses effectively evoking spikes (2→3) would be strengthened with spike-timing-dependent plasticity (STDP), whereas those arriving from a non-phase-locked area (4→3) would arrive after the target cell fires and thus be weakened. Whereas the mechanisms in (Bi-iii) would act synergistically to reinforce inphase firing between co-rippling sites within a given ripple, the mechanism in (Biv) would act across ripples to reinforce a particular network of co-rippling sites.

### Cortical ripples are associated with increased and phase-modulated single unit firing

For ripples to have a role in the communication and integration of information between distant cortical sites requires that neuronal action potentials coordinate with ripple occurrence and phase. We analyzed microarray recordings from human lateral temporal cortex granular/supragranular layers in 3 patients (STable 7) during NREM to test whether ripples modulate local single unit spiking. We detected spikes and sorted them into putative pyramidal and interneuron units based on waveform shapes and spike-timing characteristics, and verified that the units had large peak signal-to-noise ratios (PY: 9.1±3.4; IN: 5.2±2.9), had minimal interspike intervals that were <3ms (PY: 0.2±0.3%; IN: 0.3±0.6%; low percentages indicate minimal contamination by other units), and were well-isolated from one another based on the projection test (29) (PY: 95.4±86.0 SD; IN: 82.8±83.4 SD). We also detected ripples recorded by the array’s microcontacts with the average spike waveform of each unit subtracted at the times of the spikes on the unit’s channel to prevent unit spike contamination of the LFP. We found that cortical ripples were associated with increased single unit firing (Fig.5A-B). Putative pyramidal cells had a 255% increase and putative interneurons had a 297% increase in spike rates during ripples compared to baseline (randomly selected epochs in between ripples that were matched in number and duration to the ripples). Furthermore, unit firing was strongly phase-modulated by the ripples (Fig.5A), with 49% (32/66) of putative pyramidal cells and 71% (24/34) of putative interneurons having significant 70-100Hz phase modulations during local ripples (Fig.5C; post-FDR *p*<0.05, binomial test between phases within 0±π/2 vs. π±π/2, expected value=0.5, minimum 30 spikes per unit across ripples). Thus, since unit spiking is coupled to the local ripple phase, and since ripple phases are synchronized at long distances, these data suggest that ripples coordinate unit spike-timing between wide separations in the cortex, which could enable phase selection, coincidence detection, re-entrant processing, and spike-timing-dependent plasticity (Fig.6B). These basic neurophysiological processes would influence the cooperative selection of cell-assemblies between cortical areas, the essence of binding.

### Co-rippling increases putative unit-activity correlation between distant cortical sites

Our findings that co-ripples are often phase-locked across distant sites, and that unit firing is phase-locked to local ripples, together imply that unit firing is also correlated across distant sites. We were unable to test this prediction directly because we did not perform microelectrode recordings of units in multiple locations separated by more than ~5mm. Rather, as an indirect test, we used >200Hz amplitude from SEEG recordings as a proxy for unit firing (30). During waking but not NREM, this measure was more highly correlated when cortical sites were co-rippling vs. when they were not (Fig.5D, *p*=8×10^-303^, two-sided paired *t*-test). Correlation between co-rippling sites did not significantly decrement with increasing distance between the sites (Fig.5E).

As described above, co-ripple phase-lags between channels tended to be near 0 or π, especially during waking. This could indicate that co-rippling sometimes represents a state of inhibited communication, or simply that our bipolar SEEG derivations had variable relationships to local ripple generating dipoles. Indeed, the 3mm bipolar SEEG contact separation is large enough to record from multiple ripple dipoles with different orientations. In rats, ripple generators occupy ~1mm^2^ of cortical surface (5), and cellular generators are located in multiple layers separated by ~1mm (10). To test these hypotheses, we compared the correlations of the analytic amplitudes of the 200Hz highpassed signals between sites, when the sites had co-ripple phase-lags of 0±π/6 vs. π±π/6 (averaged within these ranges for each channel pair). No significant difference was found for waking or NREM (Waking: *p*=0.17; NREM: *p*=0.07, twosided paired *t*-test, *N*=2275 channel pairs), thus suggesting that the lags near 0 and π are functionally equivalent as would be expected if they are due to slight differences in electrode location relative to cortical lamina.

## Discussion

Here, using human intracortical recordings, we show that ripples couple, co-occur, and phase-synchronize across all lobes and between both hemispheres, with little decrement, even at long distances. The rate of co-rippling above chance increased exponentially with the proportion of co-rippling sites, which was correlated with stronger phase-locking. Cortical neurons increased firing during ripples and phase-locked to them, a requisite for ripples to enhance interaction via gain-modulation and coincidence detection. Enhanced long-distance interaction during co-rippling between sites was supported by increased high-frequency correlation. Ripples cooccurred between cortical sites and between the cortex and hippocampus during both spontaneous waking and NREM, and there were more co-occurrences preceding successful delayed recall. Overall, our results suggest that distributed, phase-locked cortical ripples possess the properties that may allow them to facilitate integration of the different elements comprising a particular declarative memory, or more generally, to help ‘bind’ different aspects of a mental event encoded in widespread cortical areas into a coherent representation.

Previous work showed that coupling of anterolateral temporal and parahippocampal ripples in humans increases prior to correct recall in paired-associates learning (8). We show that such coupling also occurs between hippocampal and cortical ripples. Further, we show that immediate recall with no intervening distractor, a task with identical sensory and motor stimulation but that does not require the hippocampus (12), is not associated with increased hippocampo-cortical ripple coupling. Critically, given the importance of transcortical connections between sites encoding previously unrelated elements in declarative memory, we show that cortico-cortical ripple coupling also strongly increases prior to correct recall following a delay.

Transient medial temporal inactivation disrupts both memory formation and retrieval, implying a contribution to both (31), in addition to the hippocampal role in consolidation during NREM (1–4). We found that hippocampo-cortical ripple coupling occurred spontaneously in both waking and NREM, but with different order preferences – cortex leading during waking, and hippocampus during NREM, possibly reflecting different overall flow of information during memory formation versus consolidation. The order effect during NREM was only found for hippocampal sharpwave-ripples (21), not spindle-ripples (32). Sharpwave-ripples have stronger associations with prefrontal areas supporting contextual aspects of episodic memory, and spindle-ripples with parietal areas supporting detailed autobiographical recollection. These results thus reinforce a previous suggestion that ripples make sequential contributions to consolidation (32).

A novel finding of this study is that spontaneous cortical ripples couple (within ±500ms), cooccur (≥25ms overlap), and often phase-lock (synchronize) across large areas of both hemispheres during both sleep and waking. The minimal decrement with distance in cooccurrence or phase-locking suggests either a central driving oscillator, or a population origin, such as a web of coupled oscillators (Fig.6A). The hippocampus is unlikely to act as a central driving oscillator because hippocampo-cortical coupling and co-rippling is rarer than cortico-cortical, and hippocampo-cortical phase-locking is essentially absent. A preponderance of cortico-cortical influences might be expected given that each cortical pyramidal cell receives input from thousands of other pyramids, whereas the fan-out from hippocampus (CA1 plus subiculum) to cortex is ~1:500, so each cortical pyramid gets on average less than 10 hippocampal synapses (33, 34). The thalamus is another possible source of synchronization of multiple cortical locations. However, thalamo-cortical connections are also very rare compared to cortico-cortical (35). Given the strong relationship of cortical ripples to upstates, and less strongly, to sleep spindles (11), and the role of thalamo-cortical interactions in the generation of upstates and spindles (36), it is possible that thalamo-cortical modulation via sleep waves may contribute to cortico-cortical ripple synchronization.

Conversely, a cortico-cortical network of coupled oscillators is consistent with the finding in cats that section of the corpus callosum disrupts gamma synchrony between V1 in the two hemispheres (17). Furthermore, the highly uniform ripple frequency observed across sites, cortical regions, and individual ripples suggests that local cellular and network mechanisms set the resonant oscillation frequency of each cortical module to the same 90Hz frequency, which are thereby prone to co-oscillate when excited and connected. A major role of cortico-cortical interactions in ripple co-occurrence and phase-locking is also suggested by the strong positive feedback we observed in the spread of cortico-cortical co-occurrence and in the intensity of cortico-cortical phase-locking. Specifically, cortico-cortical co-occurrence probability during waking increases from about twice chance levels for two co-occurring sites to about 20,000 times chance for co-occurrence in 25% of the sites. Similarly, during NREM, peak cortical ripple PLV between sites increases linearly with the number of additional sites co-rippling, from ~0.2 with no additional sites to ~0.9 with twelve. Note that a PLV of 1.0 would indicate perfect consistency of phase between sites over all of their co-occurring ripples.

The mechanism whereby cortico-cortical interactions could support our finding of zero-lag phase-locking at long distances, with minimal decrement up to 250mm, is unclear. Fast cortico-cortical fibers conduct ~10m/s, travelling 111mm between successive peaks of an 90Hz ripple (37). However, synchronizing projections can still be effective at multiples of the cycle time, especially with recurrent connectivity (Fig.6B). Since long-range fibers are quite rare in human cortex, whereas local connectivity and U-fibers are dense, an astronomical number of possible multi-synaptic routes exist between any two cortical locations (35). Indeed, modelling studies show that phase-locked oscillations at ~90Hz can occur in extended cortical networks, even at zero lag, provided that the neurons have multiple-path recurrent connectivity (38, 39). Such ‘polychronous’ models (40) spontaneously select paths involving multiple relays with consistent sums, and the same pair of locations can display multiple phase-lags depending on the network they are participating in, as we observed over multiple nights of sleep. Thus, although direct evidence is lacking, the most likely possibility given our findings is that ripples co-occur and phase-lock due to emergent cortico-cortical interactions.

We found that cortico-cortical co-rippling and phase-locking show consistent differences between states. Although spontaneous cortical ripple occurrence density is higher in NREM, cooccurrence density is higher in waking. Accordingly, the likelihood of multiple sites co-rippling relative to chance grows more rapidly with number of sites during waking than NREM, the increase being about 25 times larger when 25% of sites are co-rippling. Overall, the proportion of ripple peaks that occur within 25ms of each other is about five times greater in waking than NREM. Furthermore, when co-ripples phase-lock they are more likely to have near 0 phase-lag in waking than NREM. Higher activation is indicated by the ten times higher increase in >200Hz amplitude during waking vs. NREM ripples. In contrast, NREM ripples tend to recruit sequentially across the cortex (as indicated by non-zero phase-lags), with a smaller but significant increase in >200Hz amplitude. These properties may be consistent with a rapid highly synchronous recruitment of multiple cortical sites by a high activation during waking ripples, and by contrast, more sparsely activated NREM ripples. Conversely, zero-lag recruitment could be the cause rather than the effect of the greater >200Hz activation during waking.

A mean of about 16 widely-distributed cortical locations were sampled per patient. Assuming that a ripple generating module spans ~1mm^2^ of cortical surface (5), then we recorded from ~1/10000^th^ of the ripple modules. Since co-rippling is not increased at short inter-electrode distances, the proportion of recorded sites can be used as an estimate of the proportion of the cortex that is rippling. If so, then at times, as much as 25% of the cortex is co-rippling. Similarly, the usual proportion of the cortex co-rippling can be roughly estimated from the probability, given a ripple, that another site is co-rippling, ~8% (assuming that the ripple’s lead-field is comparable to its module size).

Given the strong ripple phase-modulation of cell firing we demonstrated, ripples can be expected to strongly modulate the effectiveness of arriving synaptic input, with spikes arriving on the depolarized phase being far more likely to trigger post-synaptic firing (18). Thus, there may be a strong selection for cells that project (even multi-synaptically) between co-rippling sites with a latency matching the phase-lag (possibly plus multiples of the cycle time, Fig.6Bi). Additionally, when multiple sites co-ripple, then spikes that arrive together at a third location may have greatly enhanced effectiveness (i.e., the post-synaptic site acts as a coincidence detector, Fig.6Bii) (41). Synaptic phase-selection and coincidence detection are synergistic, and would further imply positive feedback due to recurrence (Fig.6iii). Connectivity between co-rippling locations supporting in-phase interactions would be progressive because the pre- and post-synaptic activation these mechanisms induce would strengthen in-phase connections via STDP (Fig.6Biv).

Note the requisite pre-condition for this progressive strengthening of in-phase connections is that there is consistency of phase-lags across waking or NREM co-ripples of a site pair, which is what our criterion for phase-locking required. However, in addition to consistent phase-locking *across* co-ripples, phase-locking is virtually universal *within* co-ripples because the frequencies of ripples are so similar, across ripples and locations, which would enable phase-selection, coincidence detection, and re-entrance mechanisms for selection of neural networks connecting the co-rippling sites provided that the two sites are connected either mono- or poly-synaptically.

These mechanisms would function to select the members of the neural network that includes both sites, which is the core of the ‘binding-by-synchrony’ hypothesis. In neural modeling studies of such ‘polychronous’ networks (40), any two sites typically participate in multiple networks, but with a different latency for each network. Conversely, consistent phase-lag across multiple occurrences of co-ripples indicates that the two sites participate in a consistent network. In psychological binding, combinatorial associations power vast encoding spaces (e.g., letters into words) which nonetheless include some consistently associated elements (e.g., <u> usually follows <q>). Thus, the finding that phase-locking can be either consistent or inconsistent across co-ripples between 2 sites would be expected under the hypothesis that co-ripples help implement binding.

Regardless of whether phase-locking is within or between co-ripples, it is only a statistical relationship which, like ‘functional connectivity’ based on fMRI, does not prove a physical cortico-cortical interaction. For example, both within-ripple and across-ripple phase-locking could arise if the co-rippling sites had a shared abrupt depolarizing input and an intrinsic tendency to oscillate at 90Hz to such an input. However, it is much easier to construct scenarios wherein within-ripple phase-locking would arise with only shared modulations, because it only requires that the shared modulations overlap in timing by ~30ms, and no consistency across co-ripples in that timing. In contrast, across-co-ripple phase-locking would further require an extremely consistent across-co-ripple timing of the modulatory input if that were the only synchronizing influence. For example, for the site pairs in Fig.4E such modulation on a given night of NREM would need to arrive with an accuracy of about ±1ms (the full circle represents 11ms). Furthermore, this modulation latency could not be hard wired because it changes from night to night. Given the strong association of NREM ripples with upstates and waking ripples with increased very high gamma, it seems likely that shared modulation plays a role in both within-ripple and across-ripple phase-locking. However, the consistent precision of the across-co-ripple phase-locking seems to require additional synchronizing mechanisms which, given our data, we believe are likely cortico-cortical.

Through these mechanisms, cortical ripples could activate and couple with distant but related encoding areas through progressive activation of a web of cortico-cortical positive feedback connections. The strongly increased probability of co-activation and phase-locking with increasing numbers of contacts already co-activating suggests positive feedback recruitment of the most interconnected and co-activated cells into a brief synchronous ripple. When locations join the rippling network and resonate with other members, their activity level increases. The function of the ripple could thus be to act as an amplifier through resonance, leading to the rapid assembly of a network encoding the various aspects of an event, or in the case of recall or consolidation, the network encoding the various aspects of the memory.

A potential role of co-ripples in facilitating the communication and integration of information between distant cortical sites requires both the synchronized oscillations in transmembrane currents generating the co-ripples *per se,* and coordinated neuronal action potentials. We show here that single putative pyramidal and interneurons in lateral temporal cortex increase their firing during ripples in a manner that is strongly modulated by the phase of the individual ripple cycles. Therefore, the cell firing patterns necessary to support selection of activated neurons within the co-rippling locations using gain modulation and coincidence detection are in place. While we did not simultaneously record single units in multiple distant locations to directly confirm that they are co-firing, we obtained some indication that this may be the case by examining the amplitude envelope of >200Hz local field potentials. We found that the correlation between these envelopes in distant locations is greater when they are co-rippling, and furthermore, this increase in correlation does not decrease with distance, even if the sites are in different lobes or hemispheres.

The principle alternative proposed mechanism for binding relies on hierarchical convergence in multi-attribute, multisensory areas (19, 20). This mechanism seems inconsistent with the highly distributed nature of cortical processing, and poses the difficulty how to pre-represent in a small cortical module each of the combinatorial possibilities of all elements contained in all potential experiences. However, the hippocampus seems to do this, albeit temporarily, with receptive fields that combine indicators of position from multiple modalities, as well as valence, history, and context (42). Hippocampal assembly of the different elements of an event does not require integration across the hippocampus because these elements are available locally, having been pre-mixed by the entorhinal cortex and dentate gyrus. Thus, hippocampal ripples are for communication with the cortex rather than with other hippocampal sites, and indeed they are typically local within the hippocampus (32, 43); rather, their crucial co-occurrences may be with widespread cortical ripples. In this view, binding of cortical elements can occur in the absence of hippocampal input if they are previously consolidated, because cortico-cortical co-rippling and phase-locking are dependent on intracortical processes.

In summary, the characteristics of co-occurring and phase-locked cortical ripples found in the current study appear to fulfill the central requirements for a neurophysiological process that could faciliate binding-by-synchrony. First, ripples occur in all regions of the cortex in both hemispheres (required by binding because the elements of experience are encoded throughout the cortex). Second, ripples occur spontaneously throughout waking and NREM (required since binding is a ubiquitous process), and are elevated during task periods when binding may be useful (such as reassembling the components of a memory). Third, ripples last ~70ms, a duration considered in the range of ‘the psychological moment’ (44). Fourth, ripples strongly increase local firing and firing-proxy >200Hz amplitude (required for inter-areal communication). Fifth, ripples strongly phase-modulate local putative pyramidal and interneuron firing in a manner consistent with input modulation (a primary hypothesized mechanism whereby oscillations selectively amplify particular neural assemblies). Sixth, ripples strongly co-occur and phase-lock between cortical sites indicating active cortico-cortical communication (required because without communication there cannot be integration). This strong phase-locking occurs between cortical sites but not between cortex and hippocampus, indicating that trans-cortical synchrony is probably intrinsic, i.e., not projected from elsewhere. Furthermore, co-occurrence and phase-locking are minimally affected by distance (required for integration of diverse elements). Finally, >200Hz amplitude is more correlated between sites when they are co-rippling, further suggesting interareal integration of unit firing. Additional evidence from simultaneous distributed cellular level recordings as well as direct interventions in model systems will be necessary to confirm that cortical co-ripples provide the neural substrate for binding.

## Methods

### Ripple detection

Ripple detection was performed in the same way for all structures and states, based on a previously described hippocampal ripple detection method (21, 32). Requirements for inclusion and criteria for rejection were determined using an iterative process across patients, structures, and states. Data were bandpassed with a butterworth filter at 60-120Hz (forward and reverse for zero-phase shift, 6^th^ order) and the top 20% of 20ms moving root-mean-squared peaks were detected. It was further required that the maximum z-score of the analytic amplitude of the 70-100Hz bandpass (6^th^ order zero-phase shift butterworth) was greater than 3 and that there were at least 3 distinct oscillation cycles in the 120Hz lowpassed signal, determined by shifting a 40ms window in increments of 5ms across ±50ms relative to the ripple midpoint and requiring that at least 1 window have at least 3 peaks. Adjacent ripples within 25ms were merged. Ripple centers were determined as the maximum positive peak in the 70-100Hz bandpass. Ripple onsets and offsets were marked when the 70-100Hz amplitude envelope fell below 0.75 standard deviations above the mean. To reject epileptiform activities or artifacts, ripples were excluded if the absolute value of the 100Hz highpass z-score exceeded 7 or they occurred within 2s of a ≥3mV/ms LFP change. Ripples were also excluded if they fell within ±500ms of putative interictal spikes, detected as described below. To exclude events that could be coupled across channels due to epileptiform activity, we excluded ripples that coincided with a putative interictal spike on any cortical or hippocampal channel. Events that had only one prominent cycle or deflection were excluded if the largest valley-to-peak amplitude in the broadband LFP was 2.5 times greater than the third largest. For each channel, the mean ripple-locked LFP was visually examined to confirm that there were multiple prominent cycles at ripple frequency (70-100Hz), and the mean time-frequency plot was examined to confirm there was a distinct increase in power within the 70-100Hz band. In addition, multiple individual ripples in the broadband LFP and 70-100Hz bandpass from each channel were visually examined to confirm that there were multiple cycles at ripple frequency without contamination by artifacts or epileptiform activity. Channels that did not contain ripples that meet these criteria were excluded from the study.

Additional analyses were performed specifically on sharpwave-ripples and spindle-ripples detected according to our previous methods (21, 32). First, NREM data were bandpassed from 70-100Hz, and a putative ripple was detected when the 20ms moving root-mean-squared amplitude exceeded the 90^th^ percentile. Ripples were required to have at least 3 distinct oscillation cycles as described above. Broadband LFP data ±2s around ripple centers underwent 1-D wavelet (Haar and Daubechies) decomposition to detect and remove sharp transients. Ripples were classified as sharpwave-ripples based on the similarity of the peri-ripple LFP to the 400ms average biphasic waveform template created for each patient using 100-300 hand-marked hippocampal sharpwave-ripples, as quantified by the dot product between the template and the peri-ripple LFP (−100 to 300ms), and the absolute difference between the LFP value at the ripple center. Ripples were classified as spindle-ripples if the ripple center occurred during a hippocampal spindle, detected in the same way as described below for cortical spindles, on the same channel. Ripples could be classified as both sharpwave-ripples and spindle-ripples if they met both of these criteria.

### Ripple temporal relationships

Ripple cross-correlograms (peri-cortical ripple time histograms of cortical or hippocampal ripples on a different channel) were computed to assess for ripple coupling between sites. Gaussian smoothed (window=250ms, σ=50ms) ripple center counts in channel B were computed in 25ms bins within ±1500ms relative to ripple centers at *t*=0 in channel A. A null distribution was generated by shuffling (*N*=200 times) the times of ripple centers on channel B relative to the ripple centers on channel A (at *t*=0) within the ±1500 window. Pre-FDR *p*-values were calculated by comparing the observed and null distributions for each bin over ±500ms. *P*-values were then FDR-corrected for the number of channel pairs across patients multiplied by the number of bins per channel pair (45). A channel pair was determined to have a significant modulation if there were at least 3 consecutive bins each with FDR-corrected *p*<0.05 in order to minimize the possibility of false positive significance. Whether cortical ripples were leading or lagging was determined using a two-sided binomial test with expected value of 0.5, using event counts in the 500ms before vs. 500ms after *t*=0. For plots, 50ms Gaussian smoothed (σ=10ms) event counts with 50ms bins were used.

### Cortical ripple co-occurrences

Ripple co-occurrences between channel pairs were identified by finding ripples that overlapped for at least 25ms. The center of the co-occurring ripple event was determined by finding the temporal center of the ripple overlap. Conditional probabilities of ripple co-occurrence were computed by finding the probability of co-occurrence (minimum 25ms overlap) between two channels given that there was a ripple in one of the channels, separately for each channel. This was done for P(NC|NC) (both orders), P(NC|HC), and P(HC|NC). To estimate the extent of co-rippling across the cortex at any moment, the probability that a given proportion of channels was co-rippling at any time point for each patient was computed.

Observed over chance cortical ripple co-occurrence was computed as a function of the number of sites co-rippling. Ripple co-occurrence of a given number of sites required that all of those sites had at least 25ms ripple overlap. Chance was computed for each patient by randomly shuffling the ripple epochs and inter-ripple epochs of all sites 200 times and calculating the mean number of co-occurrences for each proportion of co-rippling sites (i.e., the number of channels co-rippling divided by the total number of channels, assessed for the minimum value of 2 or more channels co-rippling).

Cortical ripple co-occurrence significance for each channel pair was computed by comparing the number of observed co-occurrences (25ms minimum overlap) for each channel pair with a null co-occurrence distribution derived from shuffling ripples and inter-ripple intervals 200 times in a moving non-overlapping 5min window and counting co-occurrences.

### Ripple phase-locking analyses

To evaluate the extent to which co-occurring ripples at different sites were synchronized at different times, we used the phase-locking value (PLV), an instantaneous measure of phaselocking (27). PLV time courses were computed using the analytic angle of the Hilbert transformed 70-100Hz bandpassed (zero-phase shift) signals of each channel pair when there were at least 40 co-ripples with a minimum of 25ms overlap for each. PLVs were computed for all such co-ripples for each channel pair across a ±500ms window relative to the temporal centers of the co-ripples. A null distribution was generated by selecting 200 random times within −10 to −2 s relative to each co-ripple center. Pre-FDR p-values were determined by comparing the observed and null distributions in 5ms duration bins (averaged across five 1ms time points) within ±50ms around the co-ripple centers. These distributions for each channel pair were *across* co-ripples at each 5ms bin relative to the co-ripple center, not *within* co-ripples. These p-values were then FDR corrected across bins and channel pairs. A channel pair was considered to have significant phase-locking if it had 2 consecutive 5ms bins each with post-FDR *p*<0.05 to minimize the possibility of false positive significance. Phase-locking modulation was computed for each channel pair as the difference from the average baseline PLV within −500 to −250ms to the max PLV within ±50ms around the co-ripple center. Separate calculations were made for NREM and waking. For plotting, PLV traces were smoothed with a 10ms Gaussian window.

Phase-locking as a function of the proportion of additional sites co-rippling was computed by identifying co-ripples on the two cortical channels of interest, then sorting these events into groups based on what proportion of additional sites had a ripple that overlapped with the co-ripple by any amount of time, and computing the peak PLV and ΔPLV around the co-ripple centers. Plots of the peak PLV for channel pairs as a function of the number of additional sites co-occurring (e.g., 3 on the x-axis means 2+3=5 total sites co-rippling) are for channel pairs with significant PLV modulations.

Intra-ripple phase-locking was tested by first generating a null distribution by computing the PLV of 5 random phase-lags 1000 times. Next, the observed PLV was computed using phase-lags between the 5 peaks of each of the two ripples in a co-ripple that were closest to the co-ripple’s time center. A one-tailed *p*-value for each co-ripple was then computed as the proportion of null PLVs that were equal to or exceeded the observed PLV. The data were then FDR-corrected across all co-ripples from all cortico-cortical channel pairs from all patients.

### Ripple phase-lag analyses

The phase-lag between co-occurring cortical ripples (25ms minimum overlap) was computed for channels with significant PLV modulations (see above) by finding the circular mean (46) of the angular difference between the two 70-100Hz bandpassed ripples during their overlapping period during each ripple that co-occurred between the two channels. The mean phase-lag for a given channel pair was computed by finding the circular mean of these circular means. For each channel pair that had a significant PLV modulation, differences in ripple phase-lags by sleep night were computed for each sleep night pair where both nights had at least 30 co-ripples using the Watson-Williams multi-sample test for equal means.

### Analyses of unit spiking during ripples

Unit spiking was analyzed with respect to local ripples detected on the same contact. Ripple phases of unit spikes were determined by finding the angle of the Hilbert transform of the 70-100Hz bandpassed signal (zero-phase shift) at the times of the spikes. Unit spiking modulations during ripples were computed by comparing the number of spikes for each unit during ripples divided by the number during non-ripple epochs, i.e., epochs in between ripples, that were matched in number and duration to the ripples.

## Acknowledgements

We thank Adam Niese, Christine Smith, Christopher Gonzalez, Daniel Cleary, Eran Mukamel, Erik Kaestner, Jacob Garrett, Maxim Bazhenov, Terrence Sejnowski, and Zarek Siegel for their support. This work was supported by NIMH (1RF1MH117155-01, T32 MH020002) and ONR-MURI (N00014-16-1-2829).

## Author contributions

C.D. and E.H. designed the study. C.D., B.S., J.S., S.B., A.R., E.E., J.G., and S.C. collected the data. C.D., I.V., and X.J. analyzed the data. B.R. and S.K. provided support for the analyses. C.D. and E.H. wrote the manuscript. E.H. supervised the work.

## Data availability

The data that support the findings of this study are available at https://doi.org/10.5281/zenodo.6270017.

## Code availability

The code that support the findings of this study are available at https://doi.org/10.5281/zenodo.6270017.

