## Supplementary Information for "Widespread ripples synchronize human cortical activity during sleep, waking, and memory recall"

### SI Appendix

#### Supplementary Methods

##### Patient selection

Data from a total of 25 patients (13 female, 31±11 years old) with pharmaco-resistant epilepsy undergoing intracranial recording for seizure onset localization preceding surgical treatment were included in this study (Supplementary Table 1,7). Patients whose stereoelectroencephalography (SEEG) recordings were analyzed were only included in the study if they had no prior brain surgery; background EEG (with the exception of epileptiform transients) in the normal range; and electrodes implanted in what was eventually found to be non-lesional, non-epileptogenic cortex, as well as non-lesional, non-epileptogenic hippocampus (such areas were suspected to be part of the focus prior to implantation, or were necessary to pass through to reach suspected epileptogenic areas).

In addition, 3 of these patients were also implanted with an intracranial microelectrode (Utah Array) into cortical tissue that was suspected based on pre-surgical evaluation to be included within the region of the therapeutic resection. The implantation of the array did not affect clinical monitoring. In all patients, the tissue into which the array was implanted was later resected to gain access to the surgical focus beneath, the electrode was determined not to be implanted in epileptogenic tissue, and seizures were determined not to originate from the area where the array was implanted.

Patients were excluded from the study if they had prior brain surgery or did not have non-lesioned hippocampal and cortical channels that were not involved in the early stage of the seizure discharge and did not have frequent interictal activity or abnormal local field potentials. Based on these criteria, 25 were included in this study out of a total of 83. All patients gave fully informed written consent for their data to be used for research as monitored by the local Institutional Review Boards at Cleveland Clinic, University of California, San Diego, Oregon Health & Science University, and Partners HealthCare (including Massachusetts General Hospital),

##### Intracranial recordings

Patients were implanted with intracranial electrodes for ~7 days with continuous recordings for seizure onset localization. SEEG electrode implantation and targeting were made for purely clinical purposes. SEEG recordings were collected with a Nihon Kohden JE-120 amplifier at 1000 Hz sampling (patients S1-17), Natus Quantum LTM amplifier at 1024 Hz (patients S18-19), or Natus Xltek EMU128FS at 1000 Hz (patients S20-22). Standard clinical electrodes were 0.8mm diameter, with 10-16 2 mm long contacts at 3.5-5 mm pitch (~150 contacts/patient). In addition, patients S19 and S22 had electrodes with 0.5-2 mm long contacts at 1-2 mm pitch.

Data were also included from three patients who underwent microelectrode recordings with the Utah Array, a 10x10 microelectrode array 400 µm contact pitch and corners omitted (1-3). The insulated probe lengths are 1 or 1.5 mm long with a base of 35-75 µm tapering to uninsulated platinum 3-5 µm tip. Data were acquired at 30 kHz sampling rate with a 0.3-7.5 kHz bandpass (Blackrock Microsystems), referenced with respect to a distant wire.

During recordings the patients were reclining in their beds in the Epilepsy Monitoring Unit. During waking, typical activities were reading, thinking, looking out the window, watching TV or videos, and speaking with family, friends or hospital personnel. These distractions were prevented during performance of the memory task. We rejected periods where the recordings displayed movement artifacts, epileptiform activity, or increased slow activity which may suggest drowsiness.

**Electrophysiology pre-processing**

Offline data preprocessing was performed in MATLAB 2019b and LFPs were inspected visually using the FieldTrip toolbox (4). SEEG data were downsampled to 1000 Hz with anti-aliasing and 60 Hz notch filtered (zero-phase) with 60 Hz harmonics up to 480 Hz. Transcortical contact pairs were identified using both anatomical location (using the pre-operative MRI aligned to the post-operative CT), and physiological properties (high amplitude, coherence and inversion of spontaneous activity between contacts), and selected such that no 2 pairs shared a contact, with the exception of the 1 mm spaced hippocampal bipolar channels described above. All SEEG analyses were performed using bipolar derivations between contacts separated by 1-5 mm in cortical or hippocampal gray matter in order to ensure that activity was locally generated (5). All bandpasses, lowpasses, and highpasses were performed by applying a $3^{rd}$ order butterworth filter (MATLAB: *butter*) in the forward and reverse directions (MATLAB: *filtfilt;* $6^{th}$ order total; zero-phase shift).

**Channel selection**

Channels were excluded from analysis if they were in lesioned tissue, involved in the early stages of the seizure discharge, had frequent interictal activity, or abnormal local field potentials. From the total 2772 bipolar channels (1326 left hemisphere) in the 22 SEEG patients (S1-22), 52 hippocampal (24 left hemisphere) and 403 transcortical (180 left hemisphere) bipolar channels were selected for the analyses (Supplementary Table 1). Polarity was corrected for individual bipolar channels such that downstates were negative and upstates were positive. This was accomplished by ensuring that during NREM, negative peaks were associated with decreased, and positive peaks were associated with increased, mean ±100 ms 70-190 Hz analytic amplitude, an index of cell firing that is strongly modulated by downstates and upstates (6).

**Electrode localization**

Cortical surfaces were reconstructed from the pre-operative whole-head T1-weighted structural MR volume using the standard FreeSurfer recon-all pipeline (7). Atlas-based automated parcellation (8) was used to assign anatomical labels to regions of the cortical surface in the Destrieux atlas (9). In addition, automated segmentation was used to assign anatomical labels to each voxel of the MR volume, including identifying voxels containing hippocampal subfields (10). In order to localize the SEEG contacts, the post-implant CT volume was registered to the MR volume, in standardized 1mm isotropic FreeSurfer space, using the general registration module (11) in 3D Slicer (12). The position of each SEEG contact, in FreeSurfer coordinates, was then determined by manually annotating the centroids of electrode contact visualized in the co-registered CT volume. Each transcortical contact pair was assigned an anatomical parcel from the atlas above by ascertaining the parcel identities of the surface vertex closest to the contact-pair midpoint. Subcortical contacts were assigned an anatomical label corresponding to the plurality of voxel segmentation labels within a 2-voxel radius. Transcortical contact pair locations were registered to the fsaverage template brain for visualization by spherical morphing

(13). White-matter streamline distances between channels were computed using the 360 parcels of the HCP-MMP1.0 atlas (14), as determined by probabilistic diffusion MRI tractography (15), are population averages from (16). When two channels were in the same HCP parcel the fiber tract distance was defined as 0.

**Sleep and waking epoch selection**

Epochs included in the study did not fall within at least 1 hour of a seizure and were not contaminated with frequent interictal spikes or artifacts. NREM periods were selected from continuous overnight recordings where the delta (0.5-2 Hz) analytic amplitude from the cortical channels was persistently increased (Supplementary Table 1). Sleep epochs were confirmed by visual inspection to have normal appearing spindles, downstates, and upstates. Waking periods were selected from continuous daytime recordings that had persistently low cortical delta as well as high cortical alpha (8-12 Hz), beta (20-40 Hz), and high gamma (70-190 Hz) analytic amplitudes. When the data included electrooculography (EOG; $N$=15/17 SEEG patients S1-17), waking periods also required that the 0.5-40 Hz analytic amplitude of the EOG trace was increased. Waking epochs were required to be separated from periods of increased delta analytic amplitude by at least 30 minutes.

**Paired-associates memory task**

Paired-associates memory task data were collected from 5 patients using Presentation (Neurobehavioral Systems). Patients were instructed to memorize a series of word pairs and to recall the second word when cued with the first. A word pair was presented with text and audio (pre-recorded speech) along with an integrating image that contained semantic content of both words. After each presentation, immediate recall was assessed by cuing the first word immediately after the offset of the word pair and image. The patient then attempted to recall the word aloud. This was repeated with 8 different pairs followed by a 6 s break, and then delayed recall of the 8 pairs was assessed in the same order where the first word was cued and the patient attempted to recall the second word aloud. After 4 s the integrating image was displayed. Delayed recall was only considered successful if the second word was said aloud before the image appeared. This sequence was repeated 20 times for a total of 160 word pairs. For one patient (S18), the task was instead structured as 25 sets of 6 pairs with the integrating image shown during the encoding period but not during delayed recall, and instead during delayed recall the second word was presented after 5 s. Delayed recall was only considered successful for this patient if the second word was said aloud before it was presented after 5 s. Task performance by patient is summarized in Supplementary Table 5.

During the task, audio containing the stimulus presentations and patient voice was split and simultaneously recorded into the task presentation computer and into the clinical amplifier system synchronously with the intracranial recordings. Stimulus markers sent by the task computer were also synchronized with the intracranial recordings. Scoring of behavioral data was done offline by listening, and recall onsets were identified offline using Audacity v2.4.2 through both visual examination of the task computer recorded audio waveforms and spectrograms and confirmation through listening. The computer recorded audio was downsampled with anti-aliasing to 1000 Hz and cross-correlated with the audio recorded alongside the intracranial recordings to ensure high temporal precision of recall onsets.

Ripple occurrences and co-occurrences relative to the word cue (text and audio) preceding recall onsets prior to successful (i.e., correct response before the answer or integrating image was presented in the delayed condition) or unsuccessful (i.e., no response, incorrect response, or response after the answer or image was presented) were computed across trials, where each trial was an average across channels. Null distributions of occurrences were computed by randomly shuffling ripple or co-occurring ripple centers for each channel in each trial within ±3 s from cue onset 10 times. Observed over chance (average of null) ripple occurrence and co-occurrence histograms had 25 ms bins Gaussian-smoothed with a 100 ms window (σ=20 ms). Trial-wise ripple occurrence and co-occurrence averages in the 150-300 ms window following cue onset were computed for statistical analyses.

**Sleep spindle detection**

Spindles were detected as previously described (17). Data were bandpassed at 10-16 Hz, then absolute values were smoothed via convolution with a tapered 300 ms Tukey window. Next, median values were subtracted from each channel. Data were normalized by the median absolute deviation and spindles were detected when peaks exceeded 1 for at minimum 400 ms. Onsets and offsets were identified when these amplitudes fell below 1. Putative spindles that coincided with large increases in lower (4-8 Hz) or higher (18-25 Hz) frequency power were rejected to eliminate broadband events (e.g., artifacts) as well as theta bursts, which may extend into the lower end of the spindle range (18).

**Interictal spike detection and rejection**

Ripples and sleep graphoelements were excluded if they were within ±500 ms from putative IIS detected as follows. A high frequency score was computed by smoothing the 70-190 Hz analytic amplitude with a 20 ms boxcar function and a spike template score was generated by computing the cross-covariance with a template interictal spike. The high frequency score was weighted by 13 and the spike score was weighted by 25, and an IIS was detected when these weighted sums exceeded 130. In each patient, detected IIS and intervening epochs were visually examined from hippocampal and cortical channels (when present) to confirm high detection sensitivity and specificity.

**Time-frequency analyses**

Average time-frequency plots of the ripple event-related spectral power (ERSP) were generated from the broadband LFP using EEGLAB (19). Event-related spectral power was calculated from 1 Hz to 500 Hz with 1 Hz resolution with ripple centers at $t$=0 by computing and averaging fast Fourier transforms with Hanning window tapering. Each 1 Hz bin of the time-frequency matrix was normalized with respect to the mean power at -2000 to -1500 ms and masked with two-tailed bootstrapped significance ($N$=200) with FDR correction and $\alpha$=0.05 using -2000 to -1500 ms as baseline.

**Unit detection, classification, quality, and isolation**

Unit detection and classification was performed according to our published procedures (20-26). Data were bandpassed at 300-3000 Hz and putative unit spikes were detected when the filtered signal exceeded 5 times the estimated standard deviation of the background noise. Units were k-means clustered using the first three principal components of each spike. Overlaid spikes were examined visually and those with abnormal waveforms were excluded. Based on their waveforms, firing rates, and autocorrelograms, action potentials were clustered as arising from putative pyramidal cells or interneurons. Putative pyramidal cells had spike rates of ~0.1-0.8 Hz, long valley-to-peak and half width intervals, sharp autocorrelations, and bimodal inter-spike interval (ISI) distributions, reflecting a propensity to fire in bursts (Supplementary Table 7). By contrast, putative interneurons had spike rates of ~1-5 Hz, short valley-to-peak and half width intervals, broad autocorrelations, and a predominantly unimodal ISI distribution.

Single unit quality and isolation were confirmed according to previously established guidelines (27). Unit spikes were verified to well-exceed the noise floor based on large peak signal-to-noise ratios (PY: 9.12±3.39; IN: 5.23±2.88). Since the neuronal spiking refractory period is about 3 ms, the percent of ISIs less than 3 ms estimates the degree of single unit contamination by spikes from different units, which was very low among the units included in this study (PY: 0.21±0.34%; IN: 0.33±0.58%). Furthermore, single units detected on the same contact were highly separable according to their projection distances (28) (PY: 95.4±86.0 SD; IN: 82.8±83.4 SD). Lastly, temporal stability of unit spikes over time was confirmed based on consistency of the mean waveform shape and amplitude of each unit across recording quartiles.

**Removal of unit spikes from local field potentials**

When detecting high frequency oscillations in an LFP from a microelectrode capable of detecting unit spikes, these unit spikes may contaminate the LFP (29), leading to false positive ripple detections. Notably, lowpassing and downsampling does not eliminate the effect of the action potential on the LFP. Therefore, we implemented a modified unit spike template subtraction technique (30). Specifically, the average unit spike waveform (-500 μs to 1600 μs centered on the trough) of each unit was subtracted from the same channel's unfiltered 30kHz Utah Array data, centered on each spike. These data were then downsampled to 1 kHz and the detection of ripples was performed as described above. Extensive visual confirmation of events in the 30 kHz LFPs confirmed that this method resulted in the detection of true oscillations.

**Statistical analyses**

All statistical tests were evaluated with $\alpha$=0.05. All p-values involving multiple comparisons were FDR-corrected according to Benjamini and Hochberg (1995) (31). FDR corrections across channel pairs were done across all channels pairs from all patients included in the analysis. Box-and-whisker plots show median, mean, and interquartile range, with whiskers indicating 1.5 × interquartile range with outliers omitted. Significance of linear correlations were assessed using the significance of the correlation coefficient when there were at least 10 data points.

In temporal analyses (i.e., peri-ripple time histograms), p-values were corrected across bins and all channels. To compute p-values using randomization tests, the observed test statistic (e.g., events per bin) was compared to the null distribution of test statistics. Null distributions were randomly selected from the same epochs using 200 iterations. The p-value was computed as the proportion of null test statistics that were equal to or greater/less than the observed test statistic. Randomization tests were one-sided according to whether the observed test statistic was greater than or less than the mean of the null test statistics. Order preferences (e.g., of ripples between channels) were assessed using a two-sided binomial test with an expected probability of 0.5. To determine if a circular distribution was non-uniform, the Hodges-Ajne test was used. To determine if two circular distributions had different circular means, a Watson-Williams multi-sample test was used. To test if co-rippling between two particular cortical sites made it more likely that additional cortical sites co-rippled, we computed a $\chi^2$ test of proportions for all possible groups of three cortical channels under the null hypothesis that the co-occurrence of channel A and B has no relation to the co-occurrence of A and C. Each table was made from the number of ripples in A that co-occurred with a ripple in B only, or C only, or both, or neither, where A occurring with B or not B are the 2 rows, and A occurring with C or not C are the 2 columns.

In the paired-associates memory analysis, chance ripple occurrences (or co-occurrences) were computed for each trial (thus separately for immediate and delayed recall) as the average of 200 random shuffles per channel (pair) within ±3 s from recall onset. Only trials where there was a successful recall (of the same word pair) in both the immediate and ~60 s delayed conditions were included. Linear mixed-effects models were performed with the patient as the random effect, as follows:

$$\textit{(co-)ripples} \sim \textit{condition} + (1 \mid \textit{patient}) \qquad (1)$$

Where *(co-)ripples* was the dependent variable computed as the average ripple rate (or co-ripple rate; minimum of 25 ms overlap for each co-ripple) within 150-300 ms following the stimulus cue preceding successful recall onset across trials, with the value for each trial as the average across channels, *condition* was 1) immediate recall or chance, 2) delayed recall or chance, 3) immediate recall or delayed recall (as a ratio of observed over chance), or 4) successful recall or unsuccessful recall (as a ratio of observed over chance). Chance levels were determined separately for immediate and delayed recall. Linear mixed-effects models with the same model formula as the one described above, with the patient as a random effect, were used to evaluate channel pair co-ripple metrics (co-occurrence probabilities, PLV modulations, and proportion of pairs with significant PLV modulations) with respect to fiber tract distance.

Differences in co-ripple phase lag preferences between nights was assessed using the Watson-Williams multi-sample test for equal means. To test if unit spiking increase during ripples, a two-sample two-sided *t*-test was used to compare the spike counts during ripples vs. the spike counts during baseline periods selected in between ripples, which were matched in number and duration to the ripples. Ripple phase-modulations of spiking for each unit with at least 30 spikes during local ripples was assessed by comparing counts within $0 \pm \pi/2$ vs. $\pi \pm \pi/2$, with an expected value of 0.5. This analysis was chosen in view of animal studies indicating that cell firing during ripples tends to have maximum contrast between these intervals.

#### Supplementary Figures

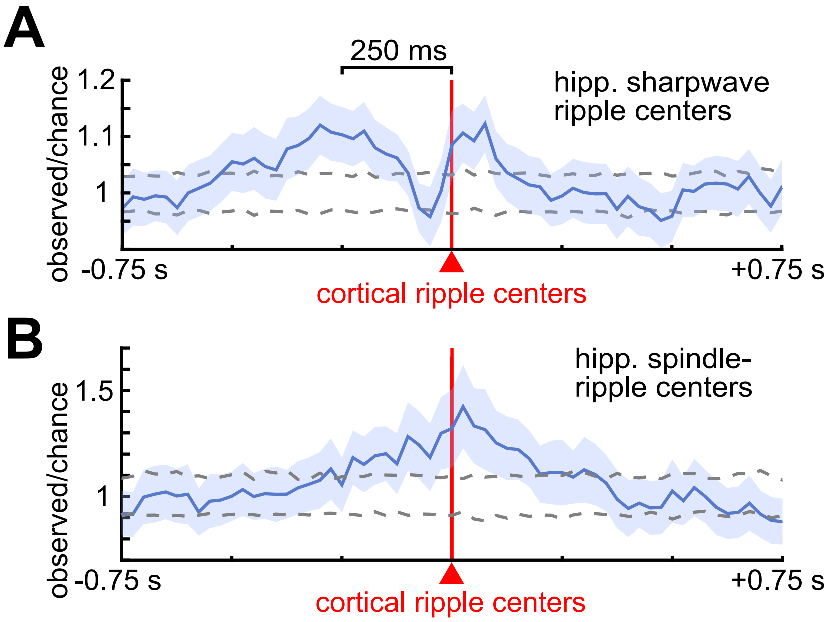


**Supplementary Fig. 1. Hippocampal sharpwave ripples precede and spindle ripples coincide with cortical ripples.** (**A-B**) Time delays from cortical ripples ($t$=0) to hippocampal sharpwave-ripples (**A**; $N$=91/461 significant channel pairs) and spindle-ripples (**B**; $N$=56/461 significant channel pairs) during NREM (post-FDR $p$<0.05, randomization test). Plots show average and SEM across significant channel pairs. Dashed curves show 99% confidence interval of the null distribution (200 shuffles/channel pair). FDR=false discovery rate correction, NREM=non-rapid eye movement sleep, SEM=standard error of the mean.

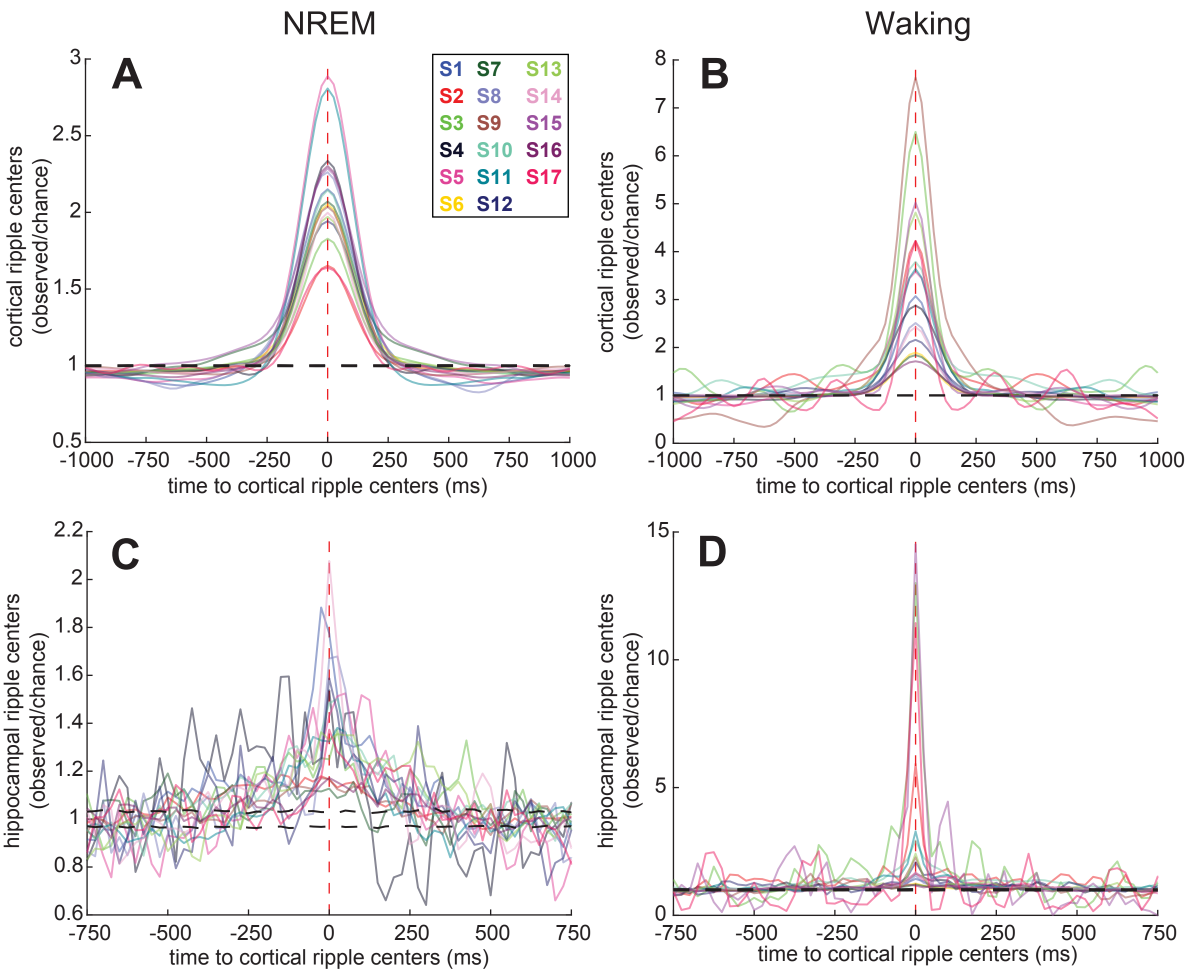


**Supplementary Fig. 2 Cortico-cortical and hippocampo-cortical ripple coupling for individual patients.** (**A-B**) Cross-correlogram of ripple times between all possible cortical sites in NREM (**A**) and waking (**B**) for individual patients (S1-17). Dashed black curves show 99% confidence interval of the null distribution (200 shuffles/channel pair). (**C-D**) Same as (A-B) except cross-correlograms of hippocampal ripple times relative to cortical ripple times. See Fig.2A-B for aggregate results.

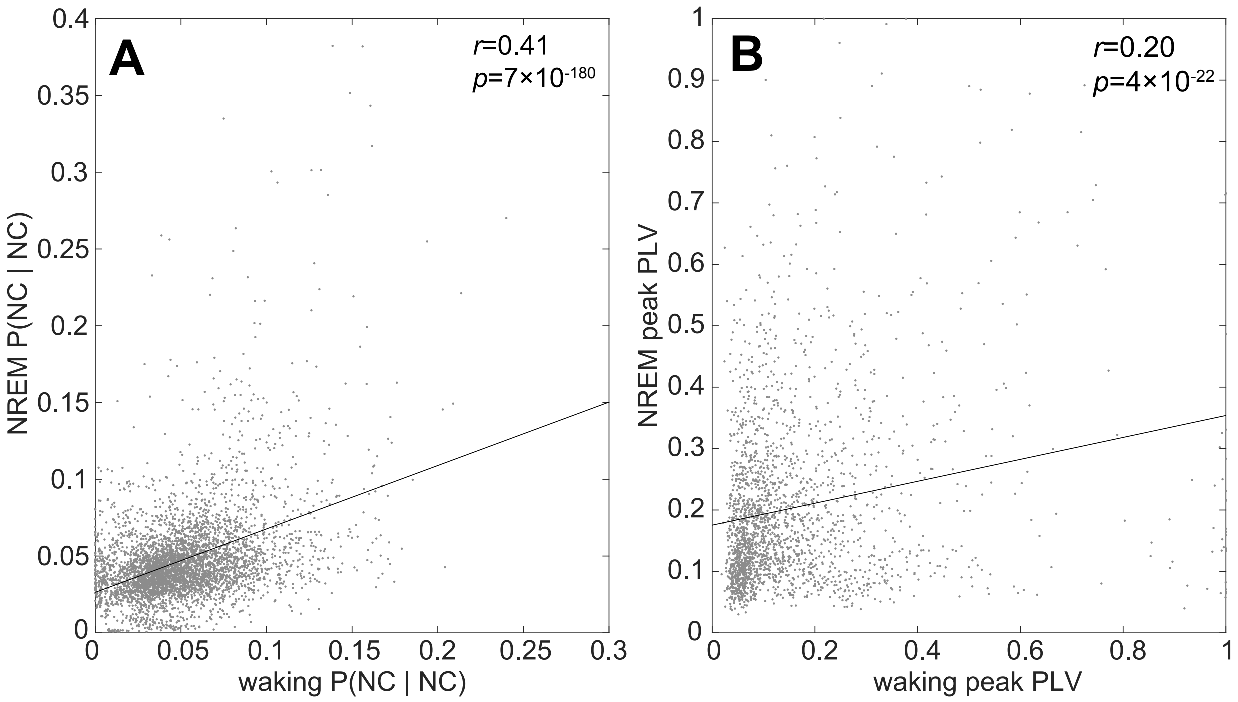


**Supplementary Fig. 3. Cortico-cortical ripple co-occurrences and phase-locking are correlated between NREM and waking.** (**A**) There was a significant correlation of the conditional co-occurrence probabilities of cortico-cortical ripples across channel pairs between NREM and waking ($r$=0.41, $p$=7×10$^{-180}$, significance of the correlation coefficient). Each point represents a channel pair. (**B**) Same as (A) except for the peak PLV during co-occurring ripples for channel pairs that had significant PLV modulations ($r$=0.20, $p$=4×10$^{-22}$). HC=hippocampus, NC=neocortex, PLV=phase-locking value.

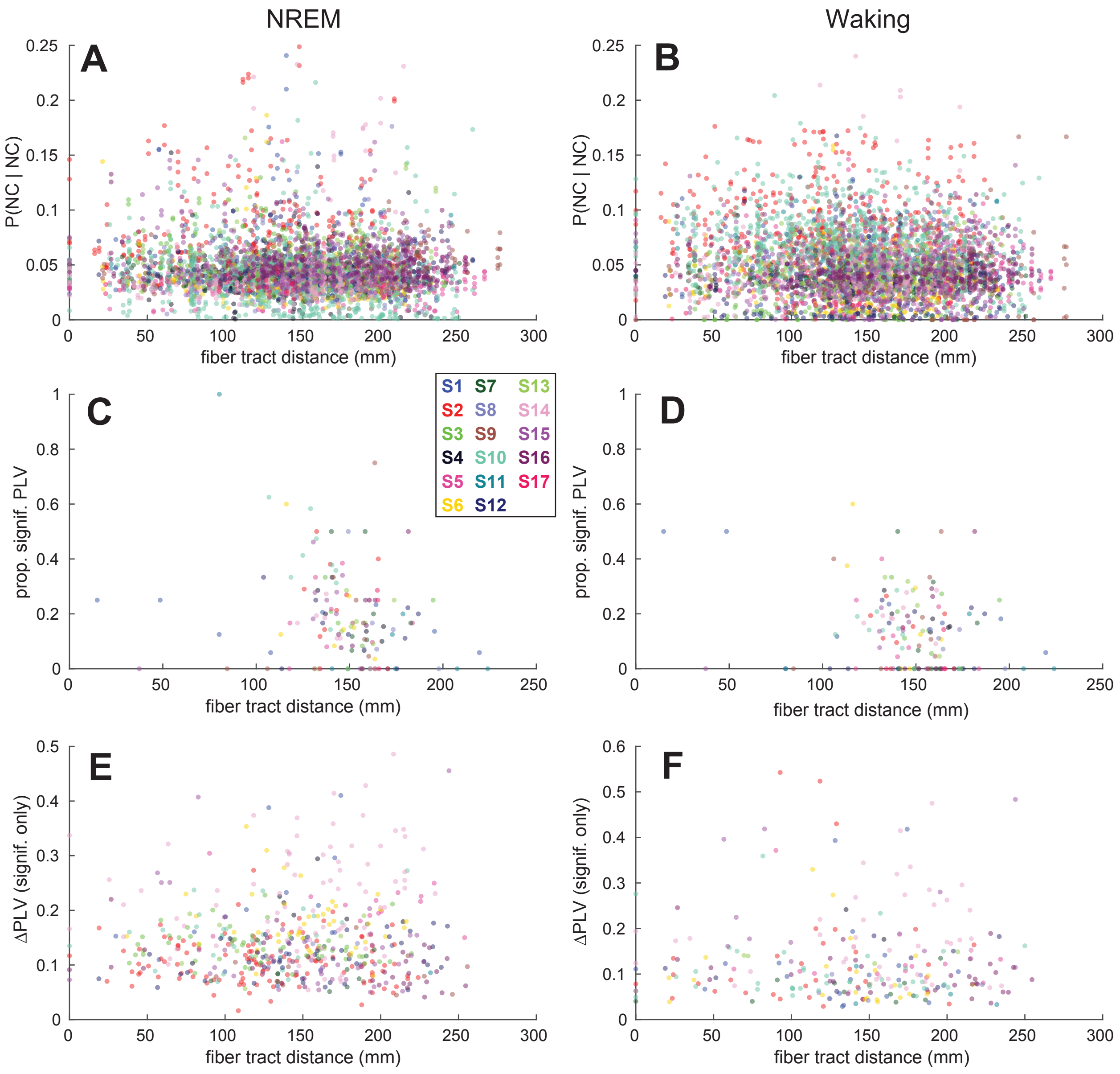


**Supplementary Fig. 4. Cortico-cortical ripple co-occurrence and phase-locking across distance for individual patients.** (**A-B**) Cortical ripple co-occurrence probabilities for each channel pair in NREM (**A**) and waking (**B**) over intervening fiber tract distance (16) for individual patients (S1-17). (**C-D**) Proportion of channel pairs with significant PLVs over distance in NREM (**C**) and waking (**D**) for individual patients. (**E-F**) ΔPLV over distance for channel pairs with significant co-ripple PLV modulations in NREM (**E**) and waking (**F**) for individual patients.

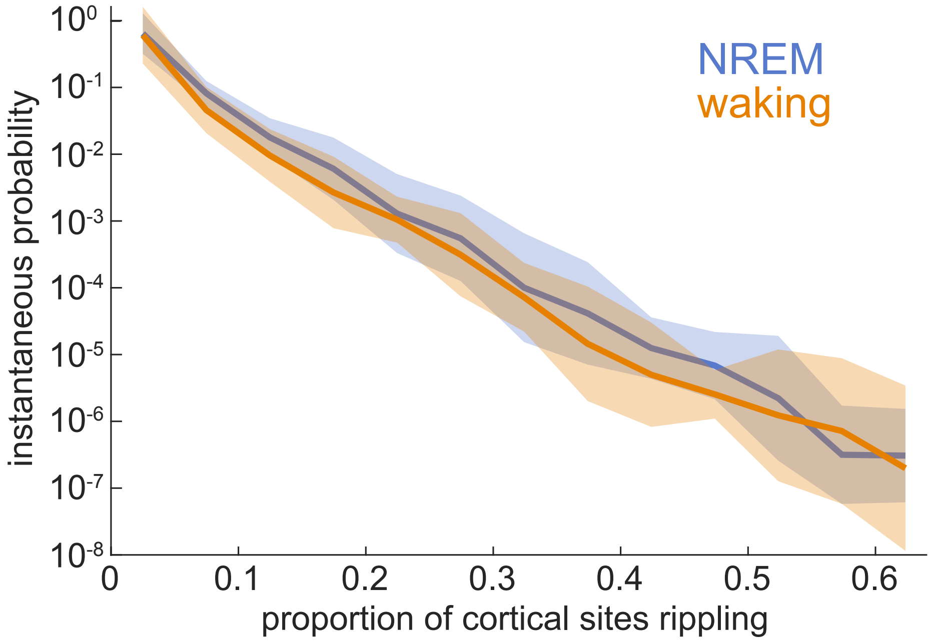


**Supplementary Fig. 5. Instantaneous probabilities of the proportions of sites rippling together.** Mean log-probability of the proportion of channels that is rippling at an arbitrary point in time during NREM or waking (*N*=273 channels from patients S1-17). Note that random co-rippling would be expected to be greater during NREM because of the greater ripple density, however, the instantaneous probabilities are similar between NREM and waking. Thus the similar curves shown here imply a relatively greater increase over chance during waking, as shown in Fig.2D. Data are binned in 0.05 increments. Error shows SEM.

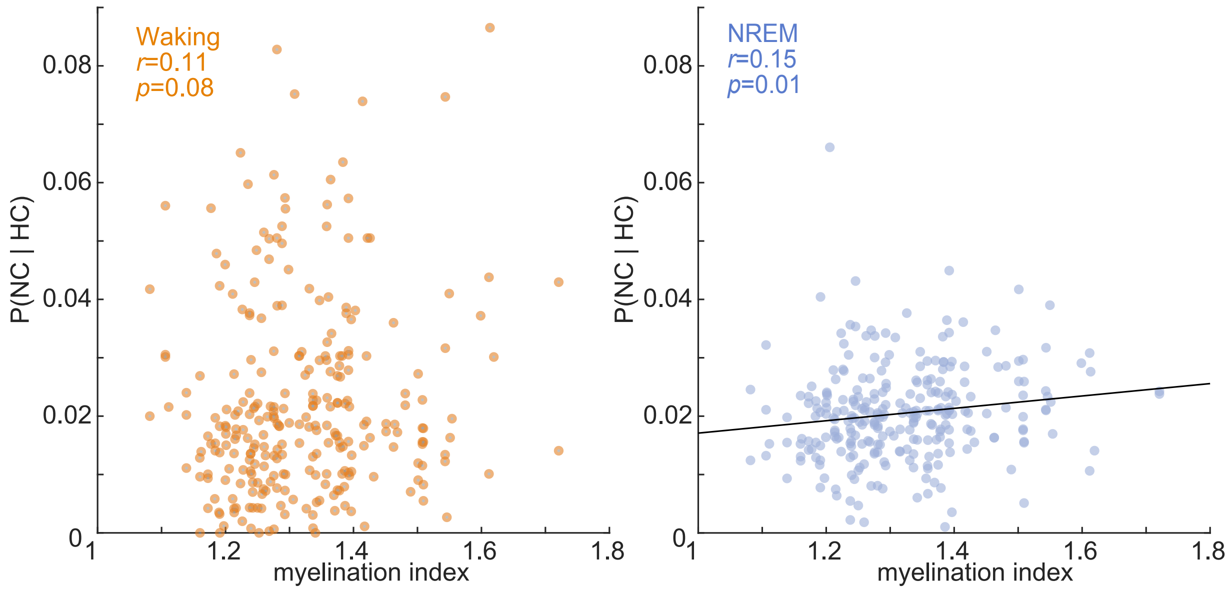


**Supplementary Fig. 6. Hippocampo-cortical ripple co-occurrence as a function of cortical myelination index.** Values are channel pair averages (each point is the average across pairs when there were multiple hippocampal channels in a patient) of the probability of a cortical ripple given a hippocampal ripple as a function of the cortical parcel myelination index (16). Co-occurrence required a minimum of 25 ms overlap.

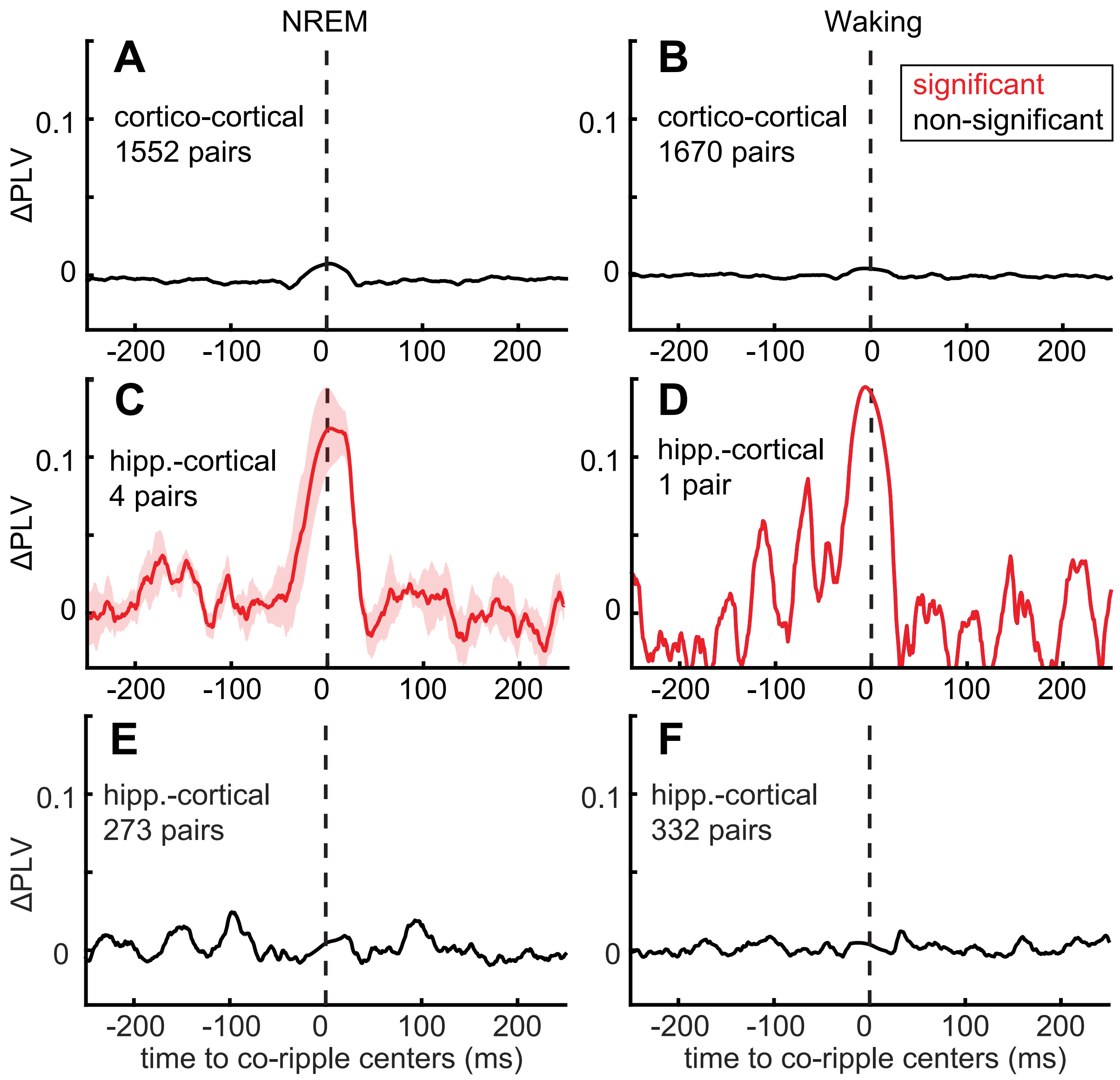


**Supplementary Fig. 7. Non-significant cortico-cortical and significant and non-significant hippocampo-cortical ripple phase-locking results. (A-B)** Average non-significant PLV time-courses locked to co-occurring ripples at $t$=0 for cortico-cortical pairs that had at least 40 co-occurring ripples during NREM (**A**) and waking (**B**). **(C-D)** Same as (A-B) except for hippocampo-cortical pairs with significant PLV responses. All significant hippocampo-cortical channel pairs localized to hippocampus proper and cortex. (**E-F)** Same as (C-D) except for non-significant PLV time-courses. Shaded error shows SEM.

#### Supplementary Tables

| Patient | Age | Sex | Hand. | No. Cort. Ch. | No. Hipp. Ch. | NREM Dur. (hr) | Waking Dur. (hr) | δ NREM / δ Waking | Paired-associates task |
|---|---|---|---|---|---|---|---|---|---|
| S1 | 20 | M | R | 18 (L) | 1 (L) | 6.1 | 49.8 | 3.98 | - |
| S2 | 58 | F | R | 22 (L) | 2 (L) | 23.7 | 23.8 | 2.35 | - |
| S3 | 42 | M | L | 16 (3 L) | 3 (2 L) | 11.4 | 19.3 | 3.39 | - |
| S4 | 18 | F | L | 15 (R) | 2 (R) | 2.7 | 2.2 | 3.02 | - |
| S5 | 20 | F | R | 18 (7 L) | 2 (R) | 5.6 | 19.3 | 2.93 | - |
| S6 | 22 | M | LR | 17 (L) | 1 (L) | 6.3 | 59.9 | 5.45 | - |
| S7 | 30 | F | R | 13 (2 L) | 1 (R) | 20.5 | 76.7 | 4.46 | - |
| S8 | 43 | F | R | 12 (L) | 2 (L) | 8.1 | 32.6 | 5.43 | - |
| S9 | 16 | M | R | 16 (4 L) | 1 (R) | 16.3 | 4.8 | 2.50 | - |
| S10 | 32 | F | R | 29 (3 L) | 3 (1 L) | 11.2 | 27.6 | 3.11 | - |
| S11 | 21 | F | L | 8 (4 L) | 3 (2 L) | 16.0 | 52.6 | 3.04 | - |
| S12 | 21 | F | R | 14 (13 L) | 1 (L) | 26.2 | 37.2 | 2.92 | - |
| S13 | 29 | F | R | 15 (6 L) | 2 (1 L) | 8.6 | 24.6 | 3.75 | - |
| S14 | 41 | F | R | 18 (R) | 1 (R) | 11.9 | 63.3 | 4.20 | - |
| S15 | 24 | M | R | 21 (9 L) | 1 (L) | 11.8 | 11.6 | 3.27 | - |
| S16 | 31 | F | R | 15 (7 L) | 1 (L) | 28.1 | 39.4 | 2.55 | - |
| S17 | 21 | M | R | 6 (L) | 1 (L) | 11.3 | 18.9 | 1.41 | - |
| S18 | 25 | M | R | 38 (R) | 0 | N/A | N/A | N/A | ✓ |
| S19 | 32 | M | L | 21 (R) | 8 (R) | N/A | N/A | N/A | ✓ |
| S20 | 34 | M | R | 22 (20 L) | 0 | N/A | N/A | N/A | ✓ |
| S21 | 38 | F | R | 21 (13 L) | 0 | N/A | N/A | N/A | ✓ |
| S22 | 23 | M | R | 28 (14 L) | 16 (8 L) | N/A | N/A | N/A | ✓ |

**Supplementary Table 1. Stereoelectroencephalography patient demographics and data characteristics.** Channels are the bipolar channels included in the analyses. δ NREM / δ Waking reports the ratio of the mean of the delta (0.5-2 Hz) analytic amplitude means across cortical channels during the NREM vs. waking epochs analyzed.

| Ripple \| Ripple | Signififcant Modulation | |
|---|---|---|
| | NREM | Waking |
| Cort-R \| Cort-R | 98.90% (3246/3282) | 97.99% (3216/3282) |
| Hipp-R \| Cort-R | 26.78% (15/56) | 64.28% (36/56) |

**Supplementary Table 2. Cortical ripple coupling with cortical or hippocampal ripples in interictal-free channels.** While the coupling analyses presented in the main text were performed between non-epileptogenic sites, during periods that did not exhibit frequent interictal spikes or artifacts (32), and with ripples that were not contaminated with interictal spikes or artifacts, additional analyses were run on a subset of hippocampal (*N*=5 channels from patients S4,6,9,17) and cortical channels (*N*=232 channels from patients S1-17) that did not have interictal spikes. These results demonstrate that cortico-cortical and hippocampo-cortical ripple coupling during both NREM and waking is present in the cases when both sites are free of interictal spikes. Numbers in parentheses are channel pair counts. *P*-values were FDR-corrected across channel pairs and bins.

| | NREM | | | Waking | | |
|---|---|---|---|---|---|---|
| Subject | Significant Modulation | Significant Sidedness | Cort-R Leading | Significant Modulation | Significant Sidedness | Cort-R Leading |
| S1 | 33.3% (6/18) | 33.3% (2/6) | 0% (0/2) | 88.9% (16/18) | 62.5% (10/16) | 100.0% (10/10) |
| S2 | 77.3% (34/44) | 50.0% (17/34) | 5.9% (1/17) | 100% (44/44) | 11.4% (5/44) | 100.0% (5/5) |
| S3 | 20.8% (10/48) | 10.0% (1/10) | 100.0% (1/1) | 83.3% (40/48) | 0% (0/40) | N/A |
| S4 | 13.3% (4/30) | 75.0% (3/4) | 0% (0/3) | 66.7% (20/30) | 25.0% (5/20) | 100.0% (5/5) |
| S5 | 22.2% (8/36) | 25% (2/8) | 50% (1/2) | 75.0% (27/36) | 66.7% (18/27) | 100.0% (18/18) |
| S6 | 0% (0/17) | N/A | N/A | 94.1% (16/17) | 25.0% (4/16) | 25.0% (1/4) |
| S7 | 15.4% (2/13) | 50.0% (1/2) | 0% (0/1) | 100% (13/13) | 46.2% (6/13) | 83.3% (5/6) |
| S8 | 16.7% (4/24) | 25.0% (1/4) | 100% (1/1) | 100% (24/24) | 25.0% (6/24) | 83.3% (5/6) |
| S9 | 56.3% (9/16) | 33.3% (3/9) | 33.3% (1/33) | 62.5% (10/16) | 20.0% (2/10) | 0% (0/2) |
| S10 | 18.4% (16/87) | 18.8% (3/16) | 66.7% (2/3) | 100% (87/87) | 21.8% (19/87) | 94.7% (18/19) |
| S11 | 45.8% (11/24) | 9.1% (1/11) | 0% (0/1) | 100% (24/24) | 12.5% (3/24) | 66.7% (2/3) |
| S12 | 14.3% (2/14) | 0% (0/2) | N/A | 64.3% (9/14) | 11.1% (1/9) | 100% (1/1) |
| S13 | 23.3% (7/30) | 57.1% (4/7) | 100% (4/4) | 80.0% (24/30) | 57.1% (4/24) | 50.0% (2/4) |
| S14 | 38.9% (7/18) | 0% (0/7) | N/A | 94.4% (17/18) | 23.5% (4/17) | 100% (4/4) |
| S15 | 0% (0/21) | N/A | N/A | 52.4% (11/21) | 0% (0/11) | N/A |
| S16 | 66.7% (10/15) | 30.0% (3/10) | 66.7% (2/3) | 86.7% (13/15) | 7.7% (1/13) | 0% (0/1) |
| S17 | 50.0% (3/6) | 33.3% (1/3) | 100% (1/1) | 100% (6/6) | 66.6% (2/6) | 0% (0/2) |

**Supplementary Table 3. Cortical ripple coupling with hippocampal ripples for individual patients.** *Significant Modulation:* Proportion of channel pairs with a significant increase in the conditional probability of a hippocampal ripple occurring in the first channel given that a cortical ripple occurred within ±500 ms in the second (one-sided randomization test, 200 shuffles, 25 ms non-overlapping bins, 3 consecutive bins each with post-FDR $p<0.05$ required for significance). *Significant Sidedness:* Those with significant modulations that had significant sidedness preference around $t=0$ (post-FDR $p<0.05$, two-sided binomial test, expected=0.5, -500 to -1 ms vs. 1 to 500 ms). *Cortical Ripple Leading:* Those with significant sidedness around 0 that had cortical ripples leading (according to counts within -500 to -1 ms vs. 1 to 500 ms). See Table 1 for aggregate results.

| Patient | % Significant | |
|---|---|---|
| | NREM | Waking |
| S1 | 9.7% | 65.3% |
| S2 | 58.5% | 60.6% |
| S3 | 12.3% | 6.8% |
| S4 | 1.5% | 0.4% |
| S5 | 1.1% | 17.9% |
| S6 | 19.7% | 46.5% |
| S7 | 4.9% | 40.2% |
| S8 | 1.8% | 96.8% |
| S9 | 23.4% | 4.5% |
| S10 | 7.6% | 80% |
| S11 | 0% | 66.1% |
| S12 | 17.3% | 5.2% |
| S13 | 5.9% | 50.8% |
| S14 | 29.8% | 74.3% |
| S15 | 17.4% | 17.8% |
| S16 | 29.5% | 22.2% |
| S17 | 0% | 4.8% |
| Average | 14.1±15.2% | 38.8±31.2% |

**Supplementary Table 4. Percent of channel-triplets where the number of triple co-occurrences significantly exceeded those that would be expected given the number double co-occurrences of the constituent channel pairs.** To test if the co-rippling between two particular cortical sites makes it more likely that other cortical sites participate, we computed a $\chi^2$ test of proportions for all possible groups of three cortical channels under the null hypothesis that the co-occurrence of channel A and B has no relation to the co-occurrence of A and C. *P*-values were FDR-corrected across channels for each patient. Error is standard deviation.

| Patient | Immediate Recall | | Delayed Recall | |
|---|---|---|---|---|
| | Accuracy | Time to Recall (s) | Accuracy | Time to Recall (s) |
| S18 | 150/150 | 1.18±0.14 | 139/150 | 1.47±0.45 |
| S19 | 157/160 | 1.35±0.13 | 66/160 | 1.82±0.66 |
| S20 | 86/104 | 0.95±0.15 | 33/104 | 2.57±0.60 |
| S21 | 158/160 | 1.07±0.23 | 103/160 | 1.76±0.66 |
| S22 | 147/160 | 1.10±0.21 | 24/160 | 2.19±0.73 |

**Supplementary Table 5. Paired-associates memory task performance for individual patients.** Values indicate successful recall counts divided by total recall cues and average time from recall cue presentation to successful recall onset. Word pair recall was considered successful when the second word was correctly stated aloud following the visual and auditory presentation of the first word. Immediate recall was assessed immediately following the presentation of the word pair to be learned and delayed recall was after ~60 s.

| Channel pair type | No. Channel Pairs | State | >40 Co-occurring Ripples | Signif. PLV / Co-occurring | Signif. PLV / All pairs |
|---|---|---|---|---|---|
| Cort ↔ Cort | 2275 | Sleep | 2106 | 26.3% (554/2106) | 24.4% (554/2275) |
| | | Waking | 1939 | 13.9% (269/1939) | 11.8% (269/2275) |
| Hipp ↔ Cort | 461 | Sleep | 277 | 1.4% (4/277) | 0.9% (4/461) |
| | | Waking | 333 | 0.3% (1/333) | 0.2% (1/461) |

**Supplementary Table 6. Significant cortico-cortical and hippocampo-cortical ripple PLVs.** Channel pairs that had ≥40 co-occurring ripples with ≥25 ms overlap were used to compute the PLVs and test for significant modulations. Signif. PLV / co-occurring reports the percent of the channels with ≥40 co-occurring ripples that had significant PLV modulations. Sig. PLV / all pairs reports the percent of all channels (no minimum co-occurrence requirement) that had significant PLVs. Results are for patients S1-17. *P*-values were computed using a randomization test (200 shuffles/channel pair, 5 ms bins within ±50 ms relative to ripple temporal centers), and were FDR-corrected for multiple bins and channel pairs, and at least 2 consecutive bins with post-FDR $p<0.05$ were required for a pair to be significant.

| Utah Array Patient Demographics | | | | | | |
|---|---|---|---|---|---|---|
| Patient | Age | Sex | Handedness | Array Implantation Location | Probe Length (mm) | NREM Duration (min) |
| U1 | 51 | F | R | Left middle temporal gyrus | 1.0 | 200 |
| U2 | 31 | M | L | Left superior temporal gyrus | 1.5 | 132 |
| U3 | 47 | M | R | Right middle temporal gyrus | 1.5 | 120 |

| Single Unit Characteristics | | | | | | | |
|---|---|---|---|---|---|---|---|
| Unit Type | No. Units | No. Spikes | Valley-to-Peak Amplitude (μV) | Spike Rate (Hz) | Valley-to-Peak Width (ms) | Half-Peak Width (ms) | Bursting Index |
| PY | 142 | 318386 | 89.57±59.46 | 0.19±0.17 | 0.49±0.06 | 0.61±0.04 | 0.05±0.03 |
| IN | 38 | 772769 | 42.26±26.11 | 1.73±1.72 | 0.30±0.05 | 0.35±0.05 | 0.01±0.02 |

**Supplementary Table 7. Single unit characteristics and patient demographics.** Errors are standard deviations. IN=putative interneuron, PY=putative pyramidal.
